# Physiological function of Flo11p domains and the particular role of amyloid core sequences of this adhesin in *Saccharomyces cerevisiae*

**DOI:** 10.1101/2021.04.01.438097

**Authors:** Clara Bouyx, Marion Schiavone, Marie-Ange Teste, Etienne Dague, Nathalie Sieczkowski, Anne Julien, Jean Marie Francois

## Abstract

Flocculins are a family of glycosylated proteins that provide yeast cells with several properties such as biofilm formation, flocculation, invasive growth or formation of velum. These proteins are similarly organised with a N-terminal (adhesion) domain, a stalk-like central B-domain with several repeats and a C-terminal sequence carrying a cell wall anchor site. They also contain amyloid β-aggregation-prone sequences whose functional role is still unclear. In this work, we show that Flo11p differs from other flocculins by the presence of unique amyloid-forming sequences, whose the number is critical in the formation of adhesion nanodomains under a physical shear force. Using a genome editing approach to identify the function of domains in Flo11p phenotypes, we show that the formation of cellular aggregates whose density increases with the number of amyloid sequences cannot be attributed to a specific domain of Flo11p. The same is true for plastic adhesion and surface hydrophobicity the intensity of which depends mainly on the abundance of Flo11p on the cell surface. In contrast, the N and C domains of Flo11p are essential for invasive growth in agar, whereas a reduction in the number of repeats of the B domain weakens this phenotype. However, expression of *FLO11* alone is not sufficient to trigger this invasion phenotype. Finally, we show that this flocculin contributes to the integrity of the cell wall.

## INTRODUCTION

The yeast cell wall is a highly dynamic structure that is not only an armour separating cell from its surrounding but that is endowed of surface properties including adherence to inert material leading to biofilm formation, cell-cell adhesion that can yield to flocculation, hydrophobicity which may result in buoyant biofilms. These properties are mediated by a variety of surface proteins called adhesin or flocculin (Orlean, 2012, Lipke *et al*., 2018). In the pathogenic *Candida albicans*, it has been reported that adhesin encoded by *ALS5* can be organised into clusters of hundreds of proteins at the cell surface leading to highly adhesion nanodomains. This 3D-organisation is triggered by an external physical shear force, such as the extension force of single molecules stretching in AFM, which can propagate across the entire cell surface at a speed of about 20 nm.min^-1^ (Alsteens *et al*., 2010). Further works by these authors demonstrated that this clustering is mediated by amyloid-core β-aggregation sequences (IVIVATT) present in Als5 protein (Garcia *et al*., 2011). In addition, Als5p has an unusual high content of β-branched aliphatic Ile, Val, and Thr that can form β-aggregates structures as predicted by the β-aggregate predictor TANGO (http://tango.crg.es/), which contributes to the cis-interactions of these proteins on the cell surface. These amyloid-like sequences and the concurrent force-induced formation of adhesion nanodomains have been recognized for other *ALS* encoding adhesins in *Candida albicans* (Otoo *et al*., 2008, Lipke *et al*., 2018). Altogether, the formation of nanodomains is an emergent property of the adhesin primary sequence, and it has been argued that the force-induced formation of these nanodomains is to strengthen cell-cell adhesion leading to the formation of robust biofilms, which promotes fungal infection (Lipke *et al*., 2018).

In a previous work aiming to investigate the impact of autolysis process on the nanomechanical properties of the cell wall of different *Saccharomyces cerevisiae* strains, we identified by Atomic Force Microscopy (AFM) the presence of abundant patches with a mean diameter of 140 nm at the surface of an industrial wine yeast strain that resembled adhesion nanodomains formed by clustering of Als5 adhesin on the surface of *C. albicans* (Schiavone *et al*., 2015). Moreover, we found that these patches resulted from the aggregation of highly mannosylated proteins since they were responsive to the binding of concanavalin A functionalized AFM tips (ConA tip), showing rupture distances that spread out over a range of 50 to 500 nm. We thus raised the hypothesis that the existence of these nanosized patches at the cell surface of this industrial yeast strain could be due to the aggregation of flocculins encoded by *FLO* genes family. The yeast *Saccharomyces cerevisiae* contains at least 5 genes encoding functional flocculins (*FLO1, FLO5, FLO9, FLO10* and *FLO11*) that share a similar architecture comprising a N-terminal sequence (A-domain), a stalk-like, repetitive and highly glycosylated B-domain and a C-terminal sequence (C-domain) that carries a glycosylphosphatidylinositol (GPI) anchoring site for covalent attachment of the protein to β (1,6)-glucans of the inner cell wall network (Dranginis *et al*., 2007, Orlean, 2012). These domains and pattern of repeats are nicely highlighted in the hydrophobic-cluster analysis (HCA) drawing shown in **Figure 1**. In addition, like *C. albicans* adhesins, the *Saccharomyces cerevisiae* flocculins contain β-aggregation sequences of 5 to 7 amino acids length that are rich in Ile, Val and Thr. The distribution of these repeats over the flocculins sequence enables to distinguish the group of Flo1p, Flo5p and Flo9p from that of Flo10p and Flo11p (**Figure 1**). In the former group, the β-aggregation-positive TANGO sequences are mainly characterized by the “T(V/I)IVI” motif that is widely distributed over the B-domain (see details in **Table S1**). Interestingly, it was shown that a TDE<u>TVIVI</u>RTP peptide containing the “TVIVI’ motif forms amyloid fibers *in vitro* (Ramsook *et al*., 2010). On the other hand, β-aggregation prone sequences are far less abundant in Flo10p and Flo11p and are located in the C-domain (Figure 1 and details in Table S1). It was furthermore reported that a 1331-residue soluble Flo11p can assemble into amyloid fibers *in vitro* most likely due to the presence of the amyloid-β-aggregation prone sequences “VVSTTV” and “VTTAVTT” at the C-terminus of Flo11p (Ramsook *et al*., 2010).

**Figure 1:**
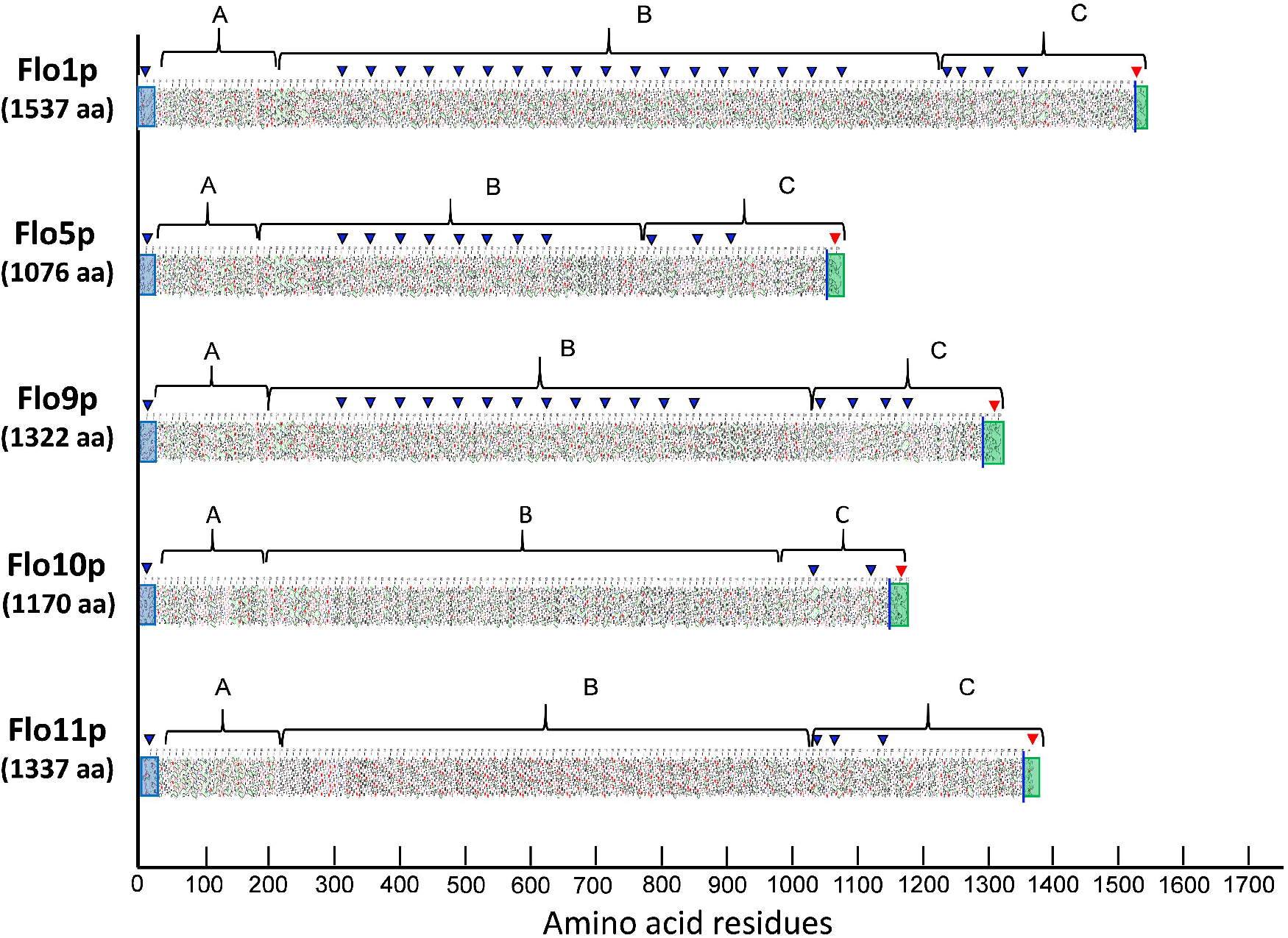
Hydrophobic-cluster analysis (HCA) of flocculins from the yeast S288C. The analysis highlights domain structure and patterns repeats. The primary structure of all Flo proteins encompasses 3 domains, namely N-terminal A domain with a secretion signal boxed in blue, a central Ser-Thr rich B-domain that carries repeat sequences with a β-aggregation potential superior to 70% marked by blue triangle, and a C-terminal domain that carries a GPI anchoring site shown by a green box and containing a highly hydrophobic sequence indicated by a red triangle.

One of the purpose of this work was therefore to study the origin and the nature of the nanoscale patches formed on the cell surface of this industrial wine yeast strain, since such abundant nanostructures have never been physically observed before in *Saccharomyces species*, despite evidence of clustering of Flo1p or Flo11p on the cell surface to account for potentiation of cell-cell aggregation under hydrodynamic shear forces and its antagonism by anti-amyloid dyes (Ramsook *et al*., 2010, Chan & Lipke, 2014, Chan *et al*., 2016). We showed in this report that these patches are adhesion nanodomains resulting from the force-induced clustering of Flo11p. We further found that the formation of these nanodomains was due to the presence of a unique sequence of 230 amino acid residues at the C-terminus of this protein, which may correspond to a sequence of 110 amino acids present in all Flo11p that is repeated two times. We then took the opportunity of this original Flo11p to investigate in greater details the importance of Flo11p domains in the various phenotypes including the formation of nanodomains that are elicited by this flocculin. This second objective was carried out by genome editing of *FLO11* using the CRISPR-Cas9 tool leading to the expression at the *FLO11* locus of Flo11p variants lacking either domain A, domain C, amyloid forming sequences, or lacking part of tandem repeats in the B-domain. Overall, this sequence-function analysis underscored that cell-cell aggregation critically depended on amyloid forming sequences, whose number potentiates this interaction leading eventually to the formation of adhesion nanodomains. However, these amyloid β-aggregation sequences have barely any effect on adherence, surface hydrophobicity and agar invasion, these later being mainly dependent on the N and the C-terminus of the Flo11p. Our results also indicate that Flo11p is necessary but not sufficient for the invasive growth phenotype and that high copy number of intragenic repeats in *FLO11* as well as high expression of this gene are not sufficient arguments to explain the flocculation and velum formation phenotype dependent on this protein.

## RESULTS

### Biophysical characterisation of the nanostructures at the cell surface of L69 strain

Our finding of abundant patches formed on the cell surface of a wine yeast L69 strain under the contact of an AFM bare tip, their persistence in autolyzed cells (Schiavone *et al*., 2015) while they were totally absent on the surface of a laboratory strain BY4741 (see **Figure S1** in supplementary data), prompted us to investigate at first the physical properties of these nanostructures by AFM using quantitative imaging (QI™) mode that enables both to image at high resolution and to quantify adhesive properties of the cell surface. **Figure 2** reports such a height and adhesion images of a single cell from L69 strain trapped in the PDMS microchamber. By zooming in a small area of the embedded cell (the white square in Fig.2A), one can clearly see that the surface of the L69 strain is rough or even dotted. This apparent roughness can be accounted for proteins that cluster together on the cell surface. From the height image at high resolution (**Fig.2B**), we estimated that these clusters have an average diameter of 100 nm with a height over the cell surface in the range of 15 −20 nm (**Fig.2C**). We then collected a total of 4096 force-distance curves from three independent cells to quantify the nanomechanical properties of the cell surface, namely adhesion forces and stiffness. Adhesion forces correspond to the retraction of the AFM tip from the surface, whereas the stiffness is the slope of the linear portion of the force versus indentation curve (see **Fig.2E**). These two physical parameters were reported as a function of the frequency of the interaction of the tip on the surface. Data of this analysis reported in **Fig 2F and 2H** showed a bimodal distribution of adhesion forces and stiffness, indicating the existence of two types of interactions. The first distribution that exhibited a mean adhesion force value of 589 pN and mean stiffness of 5.7 nN/µm likely corresponded to hydrophobic interactions (Dague *et al*., 2007). The second distribution was characterized by weaker adhesion forces (mean value of 132 pN) and higher stiffness (7.9 nN/µm), which could be attributed to cell surface proteins that unfold upon retraction of the AFM tip. To sum up, this detailed analysis by AFM showed that the nanosized patches on the cell surface of the L69 strain are adhesion nanodomains similar to those formed by Als5 at the cell surface of *C. albicans* (Alsteens *et al*., 2010). However, the observation of two different biophysical properties of these nanodomains on the surface of the L69 yeast cells raises the question of whether they can be of the same or of different biological origins.

**Figure 2:**
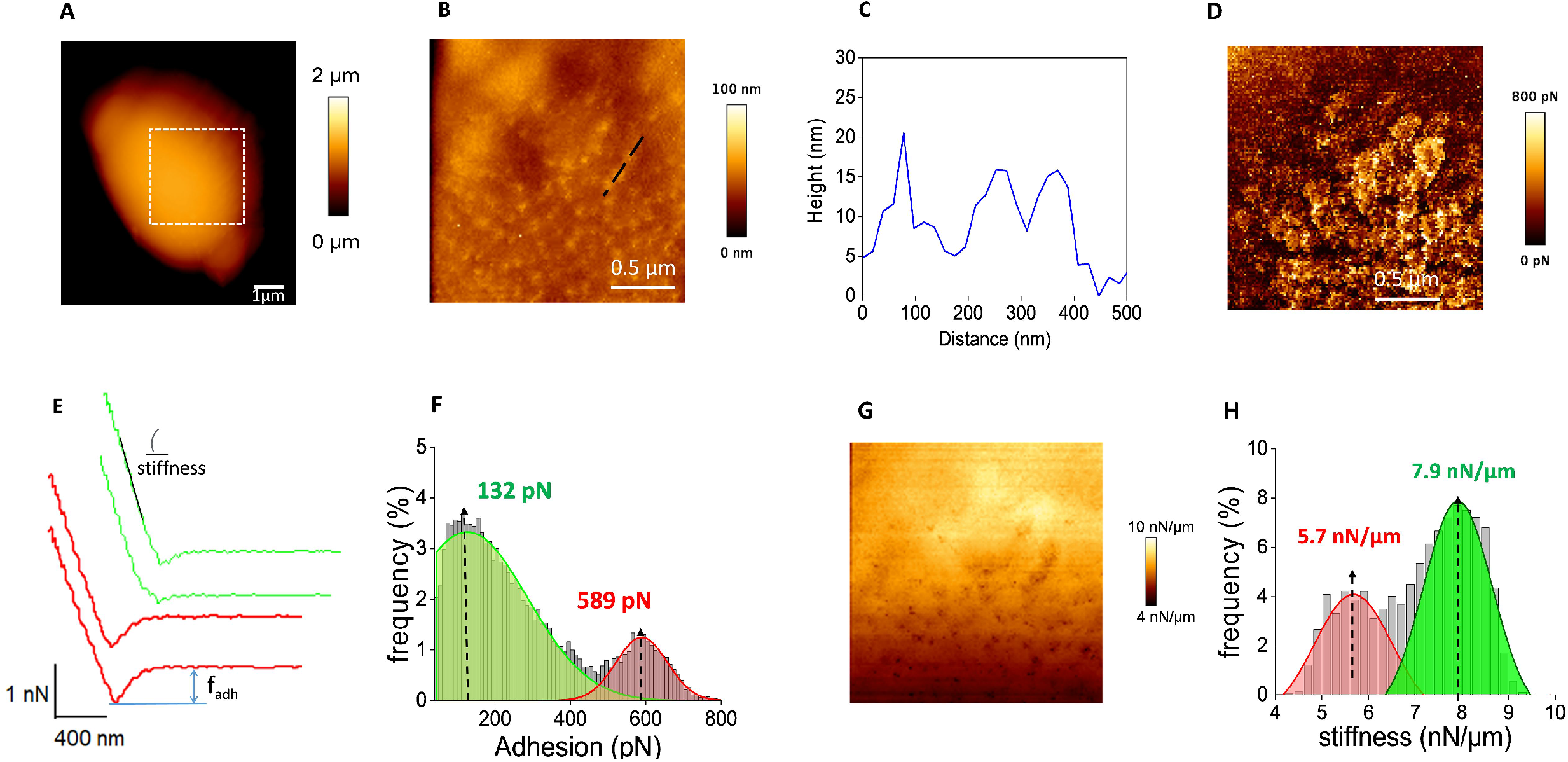
Cell surface analysis of the L69 strain using silicon-nitride (Si3N4) AFM cantilevers. AFM height image (A, B) and adhesion image (D) of a single yeast cell from L69 strain are shown. In (B) is reported an AFM contact image in which is pointed out a specific area with an arrow from which has been obtained the topography data reported in (C). In (E) are represented typical force-distance curve from high adhesive and low adhesion force and illustrated how stiffness was determined on the curves. In (F) is reported the adhesion force histogram obtained from 1024 force-distance curve recorded in QI™ mode. Cell stiffness is represented by AFM image (G) and its quantification (H) from 1024 force-distance curve recorded in QI™ mode.

### The formation of adhesion nanodomains can be inhibited by amyloid perturbants and is suppressed by deletion of *FLO11*

The model of Lipke *et al*. (Lipke *et al*., 2012, Lipke *et al*., 2018) argues that the formation of adhesion nanodomains requires amyloid-β-aggregation prone sequences in the amino acids sequence of the involved proteins. The most straightforward experiment to verify this requirement was to challenge the formation of the nanodomains in the presence of amyloid perturbants such as anti-amyloid peptide (Alsteens *et al*., 2010, Lipke *et al*., 2012) or amyloidophilic dye thioflavine S (Ramsook *et al*., 2010, Chan *et al*., 2016) that are reported to cancel the *in vivo* formation of nanodomains. Cells from L69 strain were thus treated with the amyloid disrupting peptide “VASTTVT”, which is the mutated motif of the native ‘VVSTTV’ of Flo11p from BY4741 strain (Ramsook *et al*., 2010). As shown in **Figure 3A**, a 90-min incubation with this anti-amyloid peptide resulted in a complete disappearance of the highly adhesive nanodomains. There still remained some spots of weak adhesion with values ranging from 100 to 250 pN that may likely correspond to low adhesive nanodomains as reported in Fig.2 and which can be ascribed to isolated cell surface proteins getting unfolded upon AFM tip retraction. In contrast, the two types of nanodomains were completely abolished upon treatment with 10 µM thioflavine S for 30 min (**Fig 3B**). Altogether, these data support the notion that the patches on the cell surface of L69 strain are resolved into two types of nanodomains, a low adhesive, apparently amyloid-independent, and a highly adhesive, amyloid-dependent nanodomains that are triggered by application of the mechanical force such as extension force by the retraction of AFM tip.

**Figure 3:**
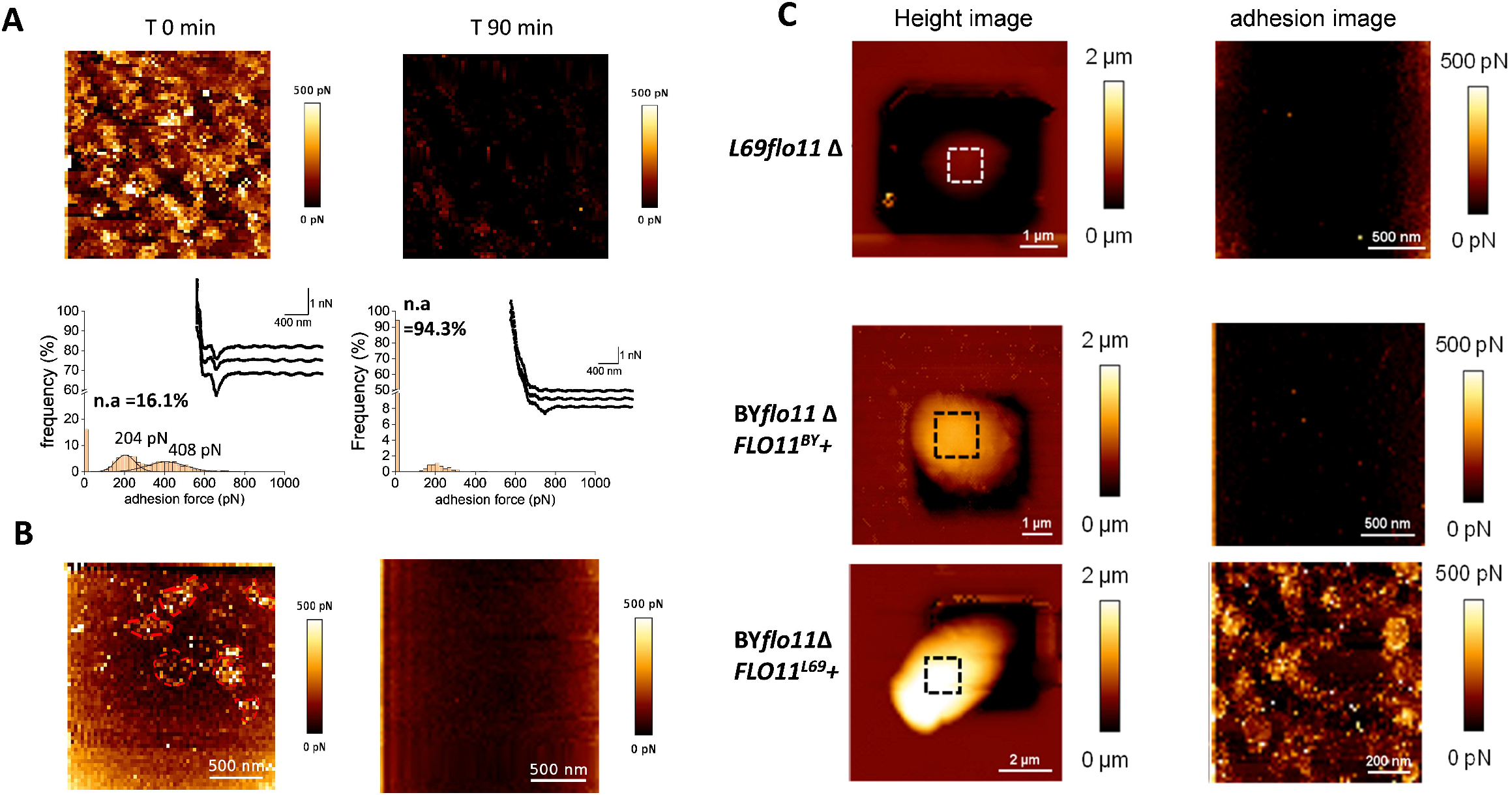
The nanosized patches at the cell surface of L69 strain are *FLO11*-dependent adhesion nanodomains that are abolished by anti-amyloïd compounds. AFM height images and adhesion images of a single cell of the *flo11*Δ mutant from L69 strain and of the laboratory BY4741 strain deleted of its endogenous *FLO11* gene (BY*flo11Δ*) and transformed with pYES 2.1 carrying *FLO11* of (BY*flo11Δ FLO11*^*BY*^+), or *FLO11* from L69 (BY*flo11Δ FLO11*^L69^+) are shown in panel A. In panel B is shown the AFM adhesion image of a cell from L69 strain before and 90 min after treatment with 5 µM of the anti-amyloid peptide VASTTV. In panel C is shown the adhesion image of a cell from a culture of L69 strain (10^7^ cells/ml) before and after 30 min of incubation with 10 µM of the anti-amyloid dye thioflavin S.

Based on our previous work showing that *FLO11* was the most highly expressed gene of the *FLO* family in strain L69 (Schiavone *et al*., 2015), and with the finding that the Flo11 protein of S288c harbours two typical amyloid core sequences “VVSTTV” and “VTTAVT” (Ramsook *et al*., 2010) near the C-terminus, we investigated the effect of deleting *FLO11* in L69 strain using the CRISPR-Cas9 toolbox (Mans *et al*., 2015). Since L69 strain is diploid (data not shown), we verified by RT-qPCR that L69*flo11Δ* did not express any *FLO11* transcript, proving that both copies of this gene was deleted (see **Figure S2A** in supplementary data). We then carried out AFM analysis in a QI^™^ mode and found that L69 defective in *FLO11* had lost the ability to form adhesion nanodomains on its surface (**Figure 3C**). This result could be an explanation of the lack of nanodomains formation in the laboratory BY4741 strain, which derives from S288c (see **Figure S1**) since *FLO11* is not expressed in this strain due to a nonsense mutation in *FLO8* that encodes a major transcriptional activator of this gene (Liu *et al*., 1996, Kobayashi *et al*., 1999). To verify this explanation, we retrieved *FLO11* gene sequence of S288c as well as that of *FLO11* from L69 strain whose genome was recently sequenced (Lallemand Inc unpublished data). Interestingly, we found that the *FLO11* gene (provisional GenBank accession number BankIt2416107 Seq1MW448340) of this industrial strain was 1.06kbp longer than that of BY4741 (*ie* 5.16 vs 4.10 kbp). Both genes were cloned into the high copy pYES2.1 plasmid under the control of *GAL1* promoter. The corresponding plasmids were used to transform BY*flo11Δ* strain (BY4741 deleted for *FLO11* by replacement with the KanMX4 cassette) and the yeast transformants were cultivated in YNGal medium that warranted its huge overexpression (see data in **Figure S2B** showing that transcript levels of *FLO11*^*BY*^ and *FLO11*^*L69*^ were respectively 2000 and 24000 fold higher than in the non-transformed BY*flo11Δ* cultivated on a galactose medium). Results undoubtedly showed that only overexpression of *FLO11* arising from L69 strain (*FLO11*^*L69*^ gene) conferred the capability of the laboratory strain to produce nanodomains **(Figure 3C)**. From these results, it can be concluded that the lack of nanodomains formation in BY4741 is not solely due the absence of *FLO11* but that the formation of these unique nanostructures requires additional features that are present in the amino acid sequence of the Flo11 protein from L69 strain.

### Comparative sequence analysis of Flo11p from various yeast strains and construction of Flo11p variants from L69 strain

Using Clustal Omega, we aligned the amino acids sequence of Flo11p from L69 strain (provisional GenBank accession number BankIt2416107 Seq1MW448340) with that of different *Saccharomyces cerevisiae* strain, including the originally genome-sequenced strain S288c (Goffeau, 1998), Σ1278b (Dowell *et al*., 2010), a strain widely studied for its remarkable properties of colony morphology, invasive and pseudohyphal growth (Reynolds & Fink, 2001, Dowell *et al*., 2010, Voordeckers *et al*., 2015) and the flor yeast strain 133d reported to form buoyancy biofilms also termed velum (Fidalgo *et al*., 2006). We found that Flo11p^L69^ is 355 and 512 amino acids longer than Flo11p^BY^ and Flo11p^Σ^, respectively, but only 92 amino acids longer than Flo11p from strain 133d (see **Figure S3** in supplementary data). This alignment also revealed that Flo11p^L69^ presented a sequence of about 230 amino acids long near the C-terminus that was totally absent in the others Flo11 proteins (see **Figure S3**, as highlighted by a red box). More remarkably, a BLAST analysis performed on all *Saccharomyces cerevisiae* strain sequenced to date revealed that this amino acids sequence was not present in Flo11p of these sequenced strains.

As already indicated in the introduction, the hydrophobicity cluster analysis tool (HCA, https://mobyle.rpbs.univ-paris-diderot.fr/cgi-bin/portal.py?form=HCA#forms::HCA) is well adapted to compare proteins having repeated sequences and high percentage of hydrophobic amino acids and hence to make emphasis on their similarity and disparity (Lo & Dranginis, 1996). As depicted in **Figure 4** (see also **Figure S4 to S7** in supplementary data for more visual details on HCA analysis of these Flo11 proteins), the Flo11 proteins of strains L69, BY4741, Σ1278b and 133d all have the same sequence architecture with an “A domain” at the N-terminus, a central B domain containing several serine/threonine (TR)-rich tandem repeats and a C-terminus carrying the GPI anchor site for attachment to the cell wall network. However, there are major differences between the Flo11p of L69 strain and that from the other yeast strains. In particular, Flo11p^69^ contains more repeats (TRs) than the proteins from BY4741 and Σ1278b (**Figure 4**), but slightly less (46 vs 49) than Flo11p from the Flor strain 133d. Interestingly, a search for the intragenic repeats in *FLO11* using EMBOSS TANDEM software (https://www.bioinformatics.nl/cgi-bin/emboss/etandem) indicated that *FLO11* from strain 133d harbored a same repeat of 81 nt that is repeated 49 times, whereas 4 different intragenic repeats with different nucleotide size were identified in the other *FLO11* genes (see details on intragenic repeats in **Table S2** in supplementary data). A second difference was in the presence in Flo11p^L69^ of two additional amyloid core sequences “VVSTTV”, which possess 75.7% β-aggregation potential (**Table 1**). A closer inspection of these amino acid residues may suggest that a sequence of about 115 amino acids length has been duplicated giving rise to an additional 230 amino acid sequence called RR2 near the C-terminus of Flo11p^L69^ (see **Figure S4 to S7** in the supplementary data). Finally, it is interesting to notice that the replacement of the ‘VVSTTV’ by the mutated ‘VASTTVT” as anti-amyloid peptide in experiments reported in **Figure 2A** dropped to 26 % β-aggregation of the sequence predicted by TANGO algorithm tool (**Table S3**).

**Figure 4:**
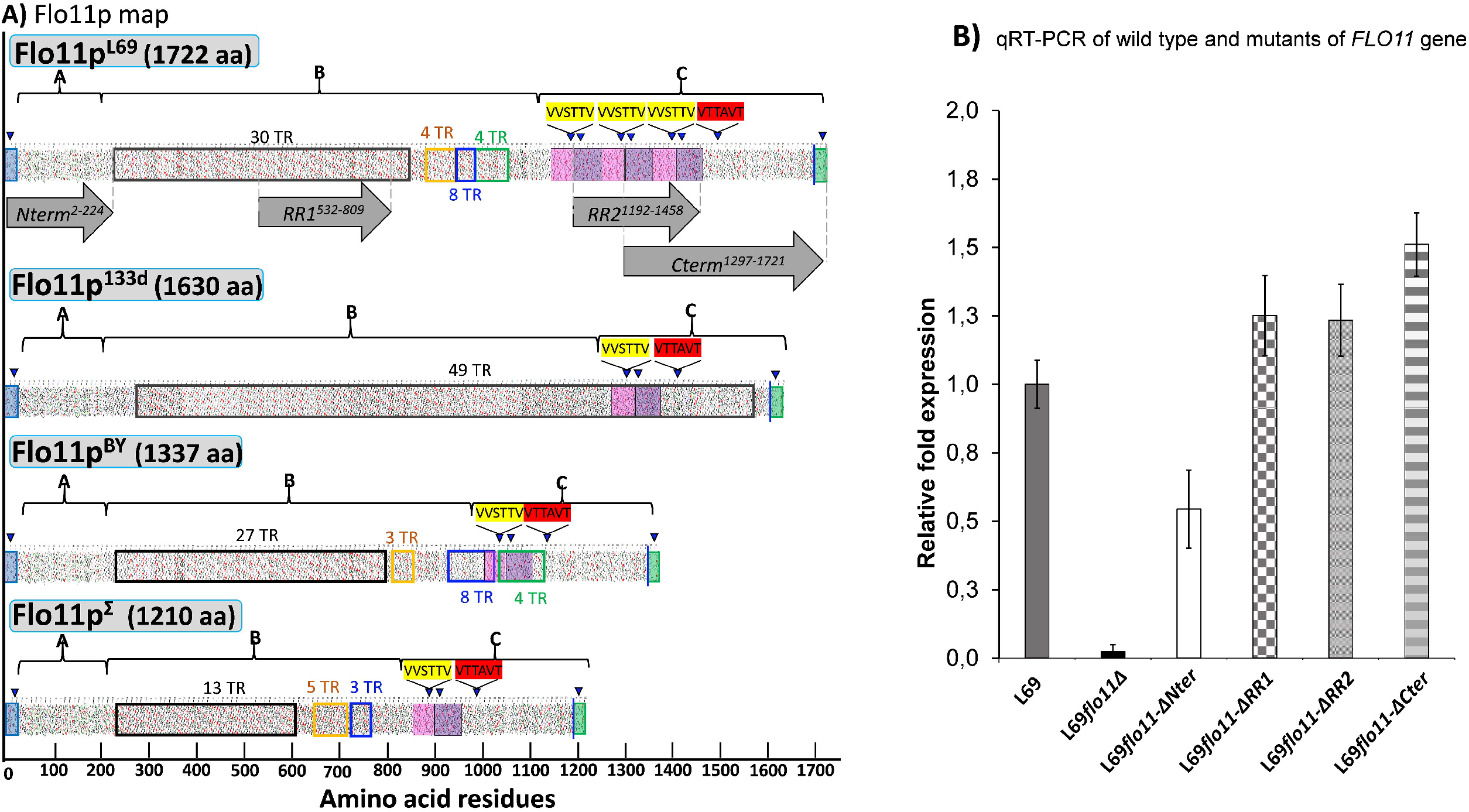
Hydrophobic-cluster analysis of the Flo11 protein from various *Saccharomyces cerevisiae* strain, and quantitative RT-PCR of wild type and mutated *FLO11* genes. In ***(A)***, is reported the HCA plots of Flo11p from wine yeast strain L69, Flor strain 133d, and laboratory strain BY4741 and Σ1278b. The three domains of the proteins as determined using various software as described in Material & Methods are highlighted by A, B and C letter. Tandem repeat domains (TR) are shown as unshaded boxes with each color addressed to a specific repeat. Blue triangles indicate regions with a β-aggregation potential superior to 30% in TANGO software and amyloid-forming sequences are indicated in yellow for ‘VVSTTV’ and red for ‘VTTAVT’. Pink and purple boxes stand for two sequence repeated three times in Flo11p ^L69^ and present only one time in the Flo11p of the other three strains. The C-terminal GPI signal is boxed in green with a blue line indicating the omega-site position (GPI signal anchorage to cell wall β-glucan). Grey arrows delimit the N-terminal, RR1, RR2 and C-terminal domains. In ***(B)***, is reported the quantitative expression levels of the different *FLO11* alleles encoding the corresponding Flo11 protein variants relative to the expression level of wild type *FLO11* in L69 strain. Samples for this qRT-PCR were taken in exponential growth phase on YN Galactose medium. Normalization of transcripts was done using *TAF10* and *UBC6* as internal reference as described in Material and Methods.

**Table 1.**
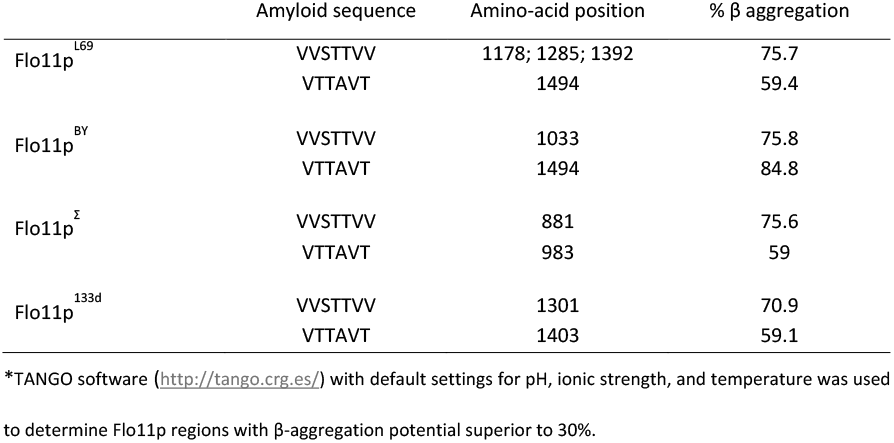
β-aggregation prone sequences in *Flo11* of various *Saccharomyces cerevisiae* strains*

| | Amyloid sequence | Amino-acid position | % $\beta$ aggregation |
| --- | --- | --- | --- |
| Flo11p<sup>L69</sup> | VVSTTVV | 1178; 1285; 1392 | 75.7 |
|  | VTTAVT | 1494 | 59.4 |
| Flo11p<sup>BY</sup> | VVSTTVV | 1033 | 75.8 |
|  | VTTAVT | 1494 | 84.8 |
| Flo11p<sup>Σ</sup> | VVSTTVV | 881 | 75.6 |
|  | VTTAVT | 983 | 59 |
| Flo11p<sup>133d</sup> | VVSTTVV | 1301 | 70.9 |
|  | VTTAVT | 1403 | 59.1 |

\*TANGO software (http://tango.crg.es/) with default settings for pH, ionic strength, and temperature was used to determine Flo11p regions with β-aggregation potential superior to 30%.

The presence of an additional RR2 region in Flo11p^L69^ raised the question of their importance in the formation of adhesion nanodomains. To answer this question, we constructed a variant of Flo11p^L69^ that was deleted of the RR2 region, as well as other variants lacking A or C-domain (deletion N and C-terminal end) by genome editing of *FLO11*^*L69*^ gene (**Figure 4A**) using the CRISPR-Cas9 technology, as described in Material & Methods. We also included a variant that bears a deletion of the repeat region RR1 which matches more than 90 % the sequence in Flo11p^BY^. The genome engineering of *FLO11* was verified by PCR to confirm that these modifications were in both copies of the gene. Moreover, expression of the genes encoding the different Flo11p^L69^ variants was verified by determining transcript levels by quantitative reverse transcription (qRT) PCR on total RNA extracted from the corresponding strains. As shown in **Figure 4B**, all *FLO11* mutants were correctly expressed and the levels of their corresponding transcripts were comparable to that of *FLO11* of the control L69 strain, with the exception that the gene coding for Flo11-ΔNter protein had an expression level about two times lower than that of the wild type gene.

### The Flo11p of strain L69 has additional amyloid β-aggregation sequences that are responsible for nanodomains formation

The cell surface of yeast strains expressing Flo11 protein variants (*ie* L69*flo11-ΔNter*, L69*flo11-ΔCter*, L69*flo11-ΔRR1* and L69*flo11-ΔRR2* strain) were imaged with AFM bare tips. As height images of individual cell trapped in the PDMS microchamber from each of these strains were similar, this argued that the expression of these Flo11p^L69^ variants had no impact on the global surface topology of yeast cell (**Fig. 5A**, left panel). However, the adhesion images at high resolution clearly revealed that yeast cells expressing the Flo11p^69^ variant lacking RR2 region (Flo11-ΔRR2) was unable to produce nanodomains under the AFM tip, while the presence of nanodomains was still recorded on the surface of yeast cell that expressed a Flo11p ^L69^ lacking either the N or the C-terminus, although the size and the morphology of these patches were different from those imaged on the surface of a L69 yeast cell expressing the wild type Flo11p (compare **Fig. 5A** with **Fig 1A**). In support of this observation, we found that nanodomains on the surface of yeast cells expressing the Flo11-ΔNter variant had a unimodal distribution of adhesion force that peaked at a maximal value of 104 pN. Moreover, these nanodomains were less stiff than those of the L69 strain expressing the wild type Flo11p (see **Figure S8** in supplementary data). On the other hand, the cell surface of L69*flo11-ΔCter* cell was characterized by needle-shaped nanostructures whose height was 3 times greater than nanodomains formed on the surface of the L69 strain. These nanodomains displayed adhesion forces that were scattered from a few pN to max 200 pN and stiffness that was roughly 30% higher than that of nanodomains from L69 strain (**see Figure S9**, in supplementary data). Finally, some tiny and disparate patches were noticeable on the surface of a L69*flo11-ΔRR1* cell that expresses a Flo11p variant in which 40% of the tandem repeats in B-domain has been removed (deletion of RR1 region, see **Figure 5A**). Typical force-distance curves recorded on these needle-shape spots showed that the adhesion forces were very weak, in the range of 100 pN (see **Figure S10** in Supplementary data), suggesting that these spots may correspond to small clusters of either Flo11 protein variant or other isolated cell wall proteins getting unfolded upon retraction of AFM tip. Taken together, these results indicated that the RR2 sequence in the Flo11 protein of L69 strain is critically important for the formation of highly adhesive nanodomains under the AFM tip, and hence suggested that a threshold number of amyloid-forming sequence is required for the formation of adhesion nanodomains.

**Figure 5:**
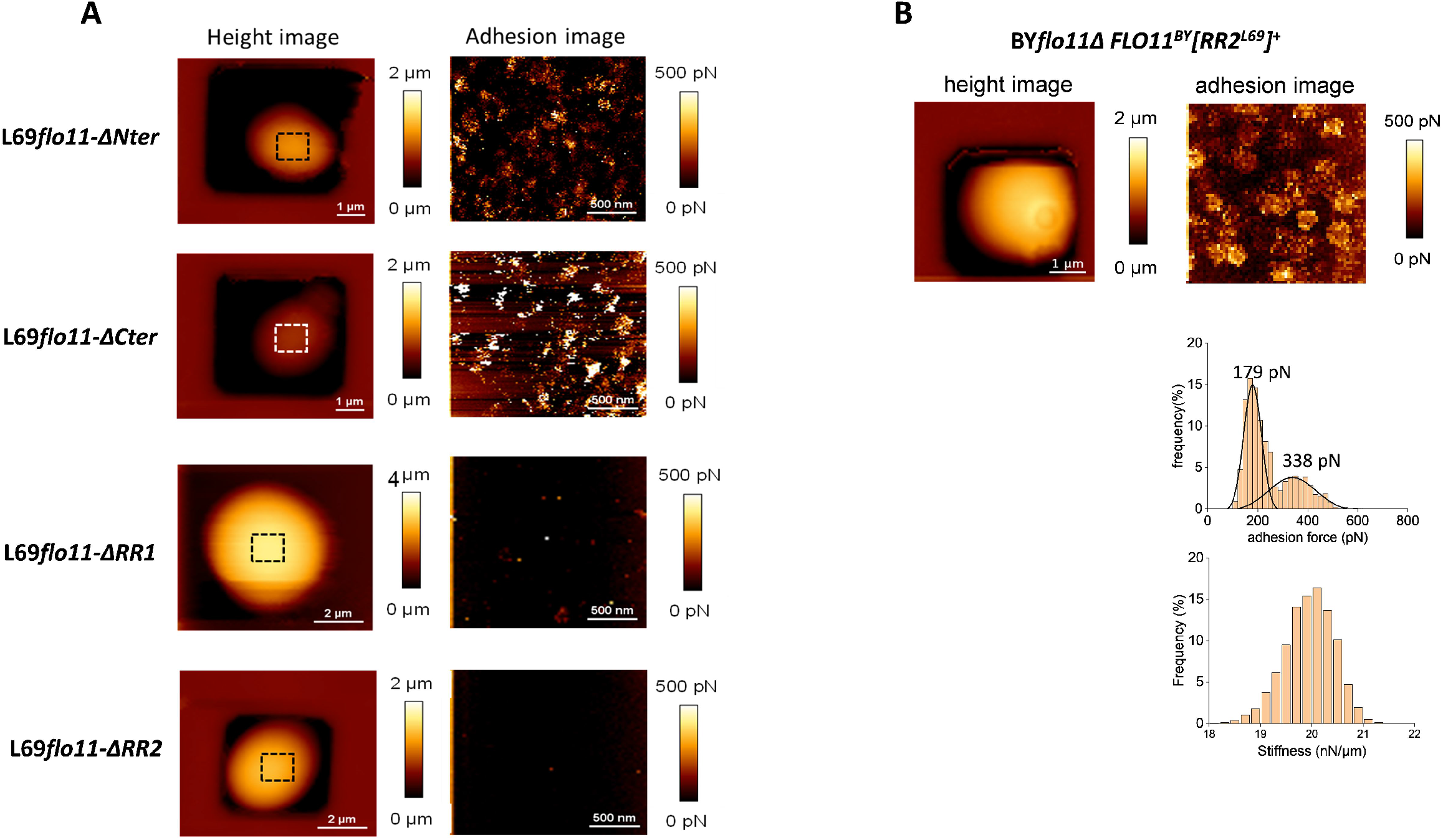
Domain in the Flo11p of L69 responsible for the production of adhesive nanodomains. In panel (A) are shown AFM height image (a) and adhesion image (b) of a single cell from L69 strain expressing Flo11p defective of the N-terminus (*flo11-ΔNter*), C-terminus (*flo11-ΔCter*) or removed from the RR1 (*flo11-ΔRR1*) or RR2 (*flo11-ΔRR2*) domain as depicted in Fig 3A. In (B) is shown AFM height and adhesion images of a single cell from BY*flo11Δ* strain transformed with pYES2.1 carrying the chimeric *FLO11*^*BY*^*[RR2]*^*L69*^ gene that corresponded to wild type *FLO11* of BY4741 in which RR2 sequence from *FLO11* of strain L69 has been inserted.

To support this assertion, we inserted the RR2 sequence of Flo11p^L69^ into the Flo11p of BY4741 and cloned this construct in pYES2.1 under the *GAL1* promoter. As shown in **Figure 5B**, low and high adhesive nanodomains were observed at the cell surface of the laboratory strain BY4741 upon ectopic overexpression of the chimeric gene encoding the Flo11^BY^-[RR2]^L69^ protein variant. The nanomechanical values of these nanodomains were however slightly different from those determined on the cell surface of the L69 strain, probably because the overall cell wall architecture of cells that overexpressed this chimeric gene is different from that of L69 cells. Nonetheless, these results confirmed the importance of the RR2 region in the force-induced formation of nanodomains, which supports the notion that a minimal threshold of amyloid core sequences within the Flo11 protein is needed to elicit this event.

### Cell-cell aggregation is enhanced by increasing the number of amyloid forming sequences

A well-established function of yeast flocculins is to promote cell-cell aggregation which can lead eventually to flocculation (Verstrepen & Klis, 2006), or to buoyant biofilm (also termed velum) that are formed by wild yeast termed ‘flor yeasts’ (Alexandre, 2013). It is considered that cell-cell aggregation proceeds by two consecutive and possibly interdependent actions, namely a cell-cell adhesion that was found to largely depend on the N-terminal domain also termed the A (adhesion) domain (Kraushaar *et al*., 2015), followed by a potentiation of these interactions by a mechanism called ‘catch bonds’. It was reported that this catch bonding is dependent on the presence of force sensitive amyloid sequences, which can lead to robust biofilms (Lipke *et al*., 2012, Lipke *et al*., 2018). Cell-cell aggregation can be easily monitored under an optical microscope by the appearance of clumps or aggregates of several cells. We quantified the intensity of this event taking into account that an aggregate must contain at least 5 cells. As shown in **Figure 6A**, both strains L69 and YSWT3a that derived from Σ1278b nicely exhibited this clumping phenotype. Deletion of *FLO11* in L69 strain almost completely abrogated the formation of aggregates whereas in the Σ1278b background strain, the size of these aggregates were dramatically reduced but surprisingly, it remained small aggregates of less than 10 cells. This residual clumping is likely a specific feature from Σ1278b background strain, which does not hamper the notion that Flo11p is critical in cell-cell interaction. This is further supported by the lack of aggregates in BY4741 as this strain does not express *FLO11* due to non-sense mutation in *FLO8* encoding its major transcriptional activator.

**Figure 6:**
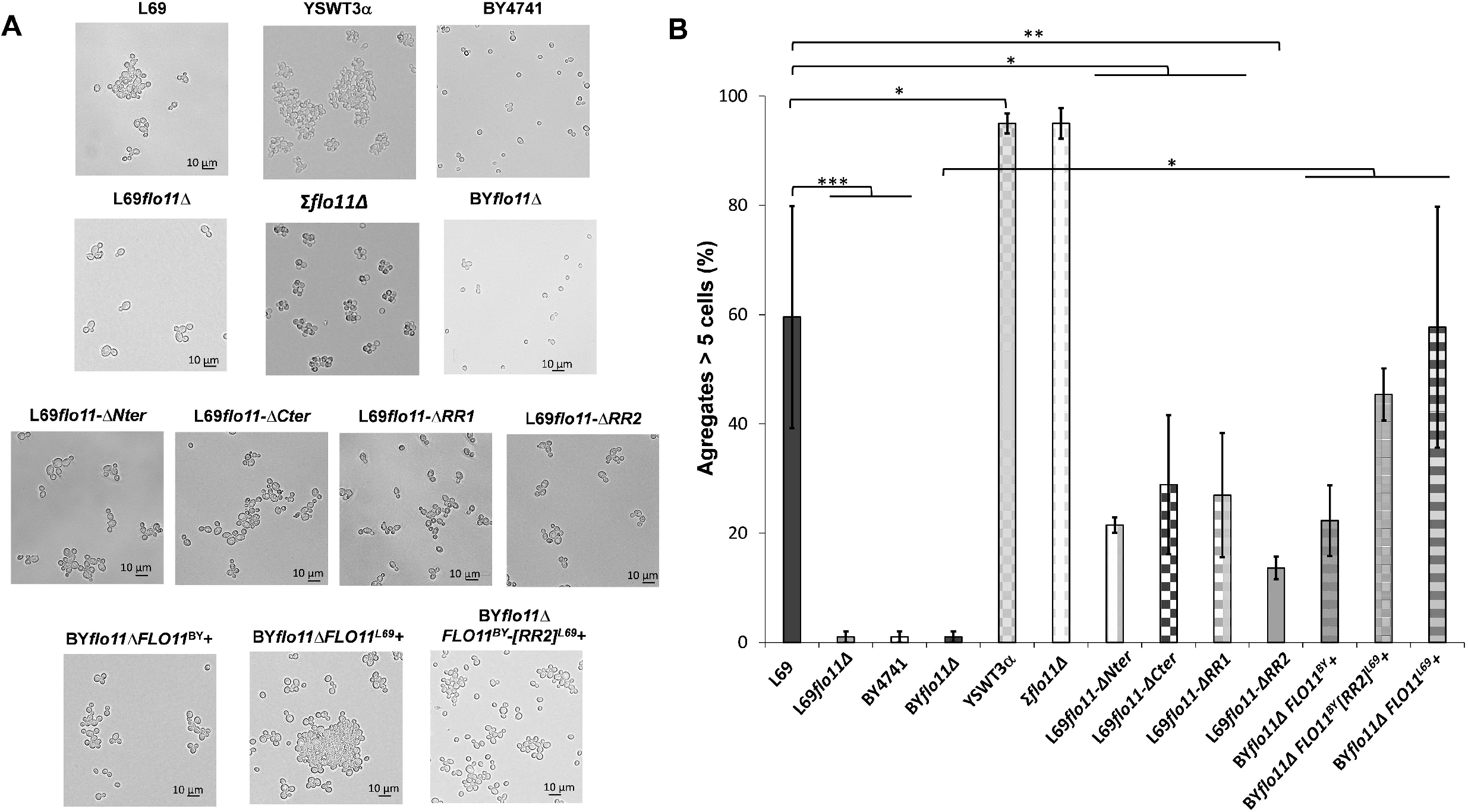
The Flo11p cell-cell aggregation is potentiated by increase number of amyloid-forming sequence. The yeast cells were cultivated as in Fig.5 but until entry in stationary phase and observed under an optical microscope. In **(A)** are shown representative photographs of cell aggregates from the various strains studied. In **(B)** is represented for each strain the percentage of aggregates that are formed by at least 5 or more cells. For each strain, more than 100 cells or aggregates were counted under the microscope. Values shown are the mean of three biological replicates and vertical bars represent standard deviations. Significant differences are denoted with asterisks (*= p-value<0.05; **= p-value≤0.01;***= p-value≤0.005).

To investigate the role of amyloid core sequence in the cell-cell aggregation, we carried out these two following experiments. On the one hand, BYflo11Δ strain that does not form aggregates was transformed with a high copy pYES2.1 plasmid carrying either its own *FLO11* gene, *FLO11* from L69 strain or a chimeric gene construct in which a DNA fragment corresponding to RR2 sequence of *FLO11*^*L69*^ was inserted into *FLO11* of BY4741 strain to yield *FLO11*^BY^-[RR2]^L69^. Following growth on galactose, cell aggregates were observed upon ectopic overexpression of these *FLO11* genes (**Fig. 6A**). However, this phenotype was much more pronounced in BYflo11 that overexpresses *FLO11*^*L69*^, reaching a percentage of cellular aggregates similar to that of strain L69. Moreover, expression of the chimeric construct *FLO11*^*BY*^*-RR2*^*L69*^ had a statistically stronger effect in aggregates formation than that of *FLO11*^*BY*^ (**Figure 6B**, see also **Figure S11** in supplementary data). Complementary to this experiment, the role of amyloid core sequences in cell-cell interaction could be assessed using amyloid perturbants. We found that aggregates formed in L69 strain or in BYΔflo11 that overexpressed *FLO11*^*L69*^ were largely disrupted upon incubation with the amyloid disruptor Thioflavine S, whereas this drug had almost no effect in L69*flo11-ΔRR2* strain that expresses a Flo11p variant lacking all amyloid-forming sequences (**Figure S11**, in supplementary data). Very interestingly, cell aggregation in YSWT3α strain was not altered by incubation with Thioflavin S, highlighting a strain background specificity for this phenotype that may be dependent on specific features in Flo11p sequence of this strain, with notably the unique presence of a 15 amino acids sequence in the A-domain that may confer stronger cell-cell adhesion property (Bruckner *et al*., 2020).

To further demonstrate the importance of amyloid core sequences in cell-cell interactions, we evaluated the effect of removing the RR2 region that bears all the 4 amyloid-forming sequence. As shown in **Figure 6**, ablation of this region in Flo11p resulted in a 75% reduction of the cell-cell interaction, and this reduction was statistically higher than that of the removal of the A-domain (Flo11-ΔNter) (**Figure 6**). The contribution of the other domains of Flo11p to this phenotype was also evaluated. The percentage of cell aggregates dropped by approximately 50% upon deletion of the C-terminal, which could be accounted by the loss of 2 out of the 4 amyloid forming sequence. A 50% reduction of cell aggregates was also observed in the strain that expresses Flo11p lacking part of the B-domain (RR1 sequence). Altogether, these results indicated that the *in vivo* efficiency of cell-cell interaction requires the full Flo11 protein.

### Contribution of various domains of Flo11p to plastic adherence and surface hydrophobicity

Using mutant strains expressing variant proteins of Flo11p^L69^, the contribution of each domain and of the amyloid core sequences on adherence and hydrophobicity properties dependent of Flo11p (Lo & Dranginis, 1996, Lo & Dranginis, 1998, Reynolds & Fink, 2001, Dranginis *et al*., 2007) could be assessed. Adherence was assayed on polystyrene surface. We found that adherence to plastic was strongly affected by pH of the culture medium, but this pH-dependency was no similar between yeast strains. While adherence to plastic of strain L69 and BY4741 was significantly reduced by raising pH from 5.0 to 8.0, it however increased in the Σ1278 derived haploid strain YSTWα. Also this property involved other cell wall proteins since deletion of *FLO11* significantly reduced but not annihilate adherence of cells to the polystyrene (**Figure 7**). Moreover, the moderate adherence of BY4741 cell to plastic at pH 5.0 must be due to cell wall proteins other than Flo11p, since this gene is not expressed in this strain due to a non-sense mutation in *FLO8* encoding its transcriptional activator (Liu *et al*., 1996). However, the critical role of Flo11p is in adherence to inert surface was clearly illustrated in BYΔflo11 strains that overexpressed *FLO11* gene (**Figure 7**). However, it is worth noticing that cells overexpressing *FLO11*^*L69*^ had higher adherence to plastic (*p-value* < 0.005) than those expressing *FLO11*^*BY*^, whether the assay was carried out at pH 5.0 or 8.0, although the intensity of this phenotype was lower at pH 8.0. This difference was probably due to the strong expression of *FLO11*^*L69*^ rather than to additional presence of amyloid sequences in Flo11p^69^ since overexpression of the chimeric construct *FLO11*^*BY*^*-RR2*^*L69*^ in BY*flo11Δ* cells gave rise to similar adherence as overexpression of *FLO11*^*BY*^. Further analysis of the domains of Flo11p that are implied in adherence to polystyrene showed that the C-terminal domain contributed the most to this property at pH 5.0, whereas at pH 8.0, this contribution relied almost exclusively on the N-terminal domain of this protein. This result was in part different from the data of Krausahaar *et al*. (Kraushaar *et al*., 2015) who showed that only the A-domain (N-ter) of Flo11p is required for cell adhesion to polystyrene.

**Figure 7:**
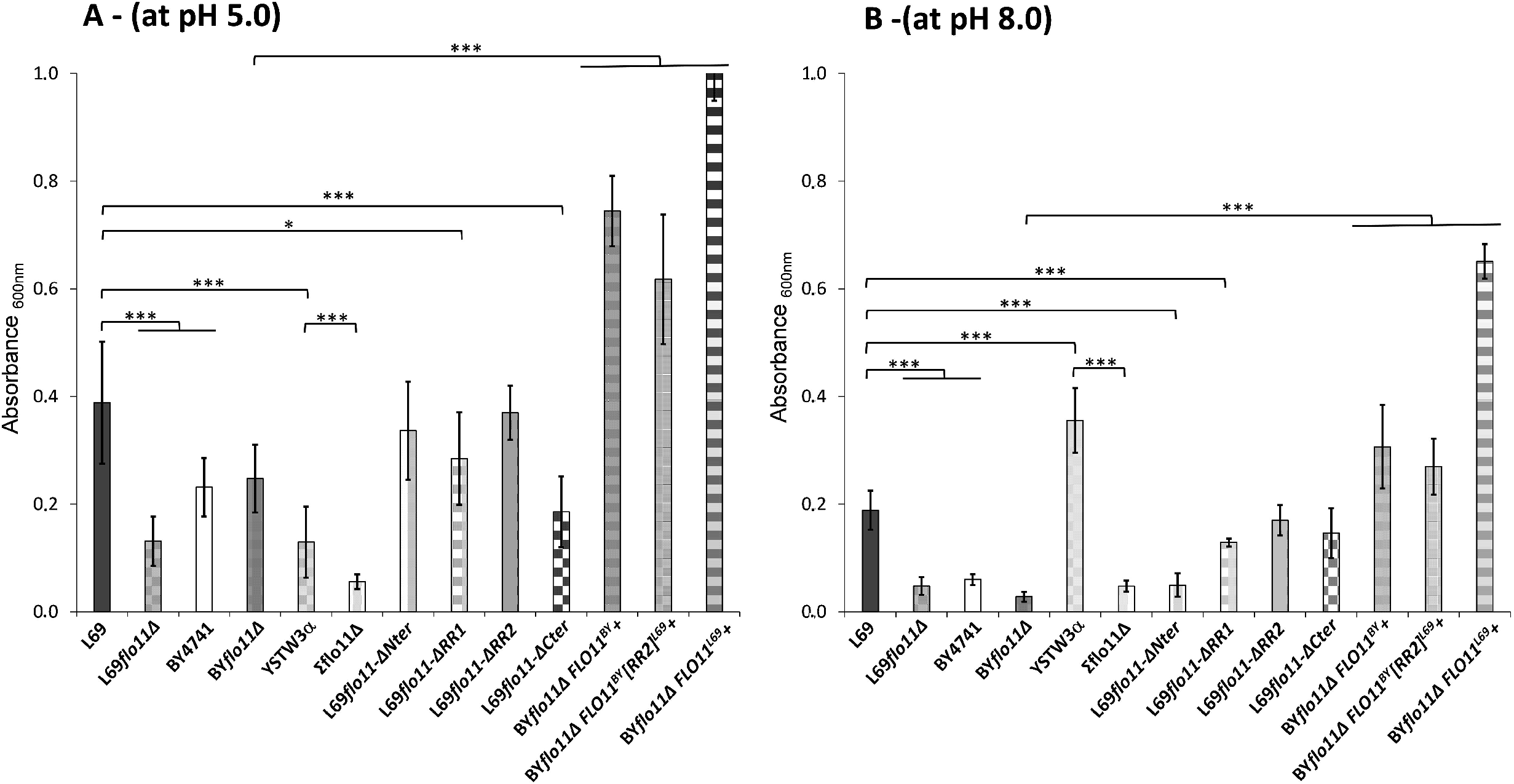
Partial Flo11p and pH-dependencies of adherence to plastic. Adherence of yeast cells was carried out a 96-well polystyrene plate as described in Methods. The data are the mean of three replicates measurements ± standard deviation. Significant differences are denoted with asterisks (*= p-value<0.05; **= p-value≤0.01;***= p-value ≤ 0.005).

Results on surface hydrophobicity, which was evaluated as the percentage of yeast cell partitioned in the octane phase, showed that each domain of Flo11p contributed to the intensity of this phenotype, with the N-terminal carrying the higher contribution and the C-terminal the lowest. However, this phenotype is mainly dependent on the amount of Flo11p exposed at the cell surface as shown by the significant increase in hydrophobicity of the cells that overexpressed *FLO11*, regardless the strain that originated this gene (**Figure 8**). Since surface hydrophobicity is critical in the formation of buoyant biofilm that is strictly dependent on Flo11p (Fidalgo *et al*., 2006), we also examined whether L69 strain, which is a wine yeast had this ability to form this air-liquid biofilm also termed velum (Alexandre, 2013). Contrary to expectation, neither L69 strains nor strain bearing any of the Flo11p variant as well as BY4741 were able to form velum, whereas this phenotype was nicely visualized in the flor strain A9 as described in ((Zara *et al*., 2005), **Figure S12** in supplementary data). Taking into account that the number of repeats in the B domain is of paramount importance in eliciting velum formation (Fidalgo et al., 2006), the absence of this phenotype in strain L69 cannot be attributed to the lack of domain repeats, since there are almost as many in Flo11p of strain L69 as in the protein of strain 133d, nor to the expression of *FLO11*, since it is particularly strong in strain L69.

**Figure 8:**
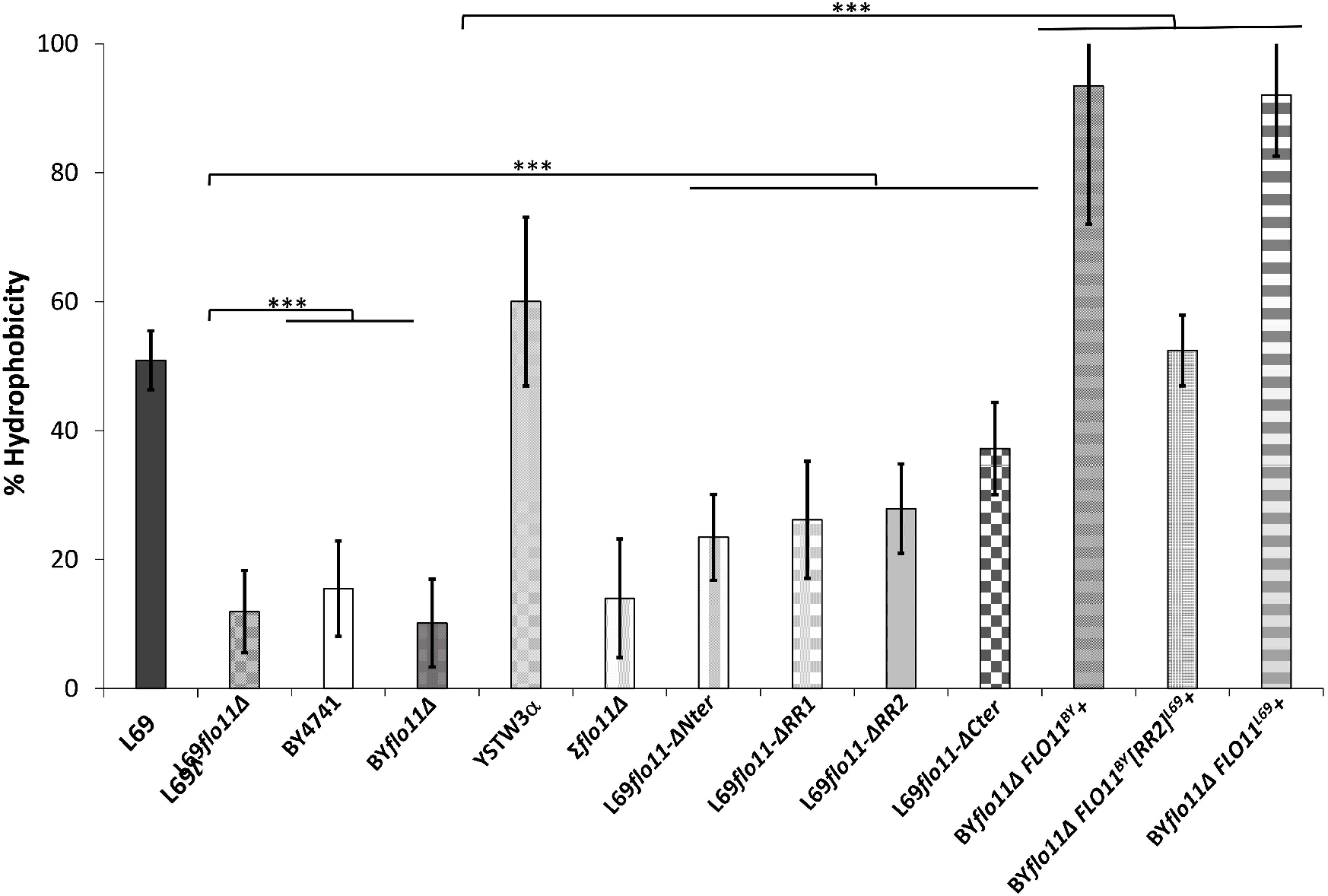
Surface hydrophobicity mainly depends on abundance of Flo11p. Surface hydrophobicity corresponded to the percentage of cells partitioning in the octane layer, as described in Methods. The data are the mean of three biological replicates and vertical bars represent standard deviations. Significant differences are denoted with asterisks (*= p-value<0.05; **= p-value≤0.01;***= p-value≤0.005).

### The Flo11p-dependent Invasive growth phenotype requires N and C-terminal of the protein, but its elicitation and intensity requests other factors that are defective in S288c background strain

It is well established that *FLO11* gene is required for pseudohyphal development in diploids and for invasive growth in haploids strains of *S. cerevisiae* (Gancedo, 2001), although the invasive phenotype can be provoked in diploid strains upon overexpression of this gene (Lo & Dranginis, 1998). This phenotype is commonly assessed by agar invasion assay that consists in cultivating yeast cells on agar plates containing rich or synthetic sugar medium for several days and then examining those cells that remained sticky on the agar plates after extensive washing under a stream of water (Roberts & Fink, 1994). Applying this assay to L69 strain, we found that this strain exhibited massive invasion in agar on both rich (YPD) and synthetic (YNGal) sugar medium (**Figure 9**). The finding that invasiveness on agar occurred in this strain, which is diploid may be attributed to the relatively high expression of *FLO11* measured in this strain (Schiavone *et al*., 2015). As expected, agar invasion was noticed for the haploid YSWT3α strain, which is derived from Σ1278b (Dowell *et al*., 2010), whereas it was absent in BY4741 strain, likely because *FLO11* is weakly expressed in this strain (Liu *et al*., 1996). Accordingly, invasion phenotype was lost upon deletion of *FLO11* in L69 and YSTW3α strains, which confirmed the critical function of this gene for this phenotype (Lo & Dranginis, 1998). As already noticed by Guo *et al*. (Guo *et al*., 2000), invasion was less pronounced on agar plates made with a galactose medium (see **Figure S12** in supplementary data). In addition we found that cells remaining the more sticky after washing are those at the periphery of the spot when the invasion experiment was carried out in a rich sugar medium, whereas cells at the heart of the spot showed the most invasiveness in a synthetic medium (**Figure 8** and see also **Figure 11** in supplementary data). More surprisingly, the diploid Σ1278b and isogenic derivative haploid YSTW3α were unable to elicit an invasion growth phenotype in a synthetic glucose medium, whereas this capacity was still very effective in L69 strain (see **Figure S12** in supplementary data).

**Figure 9:**
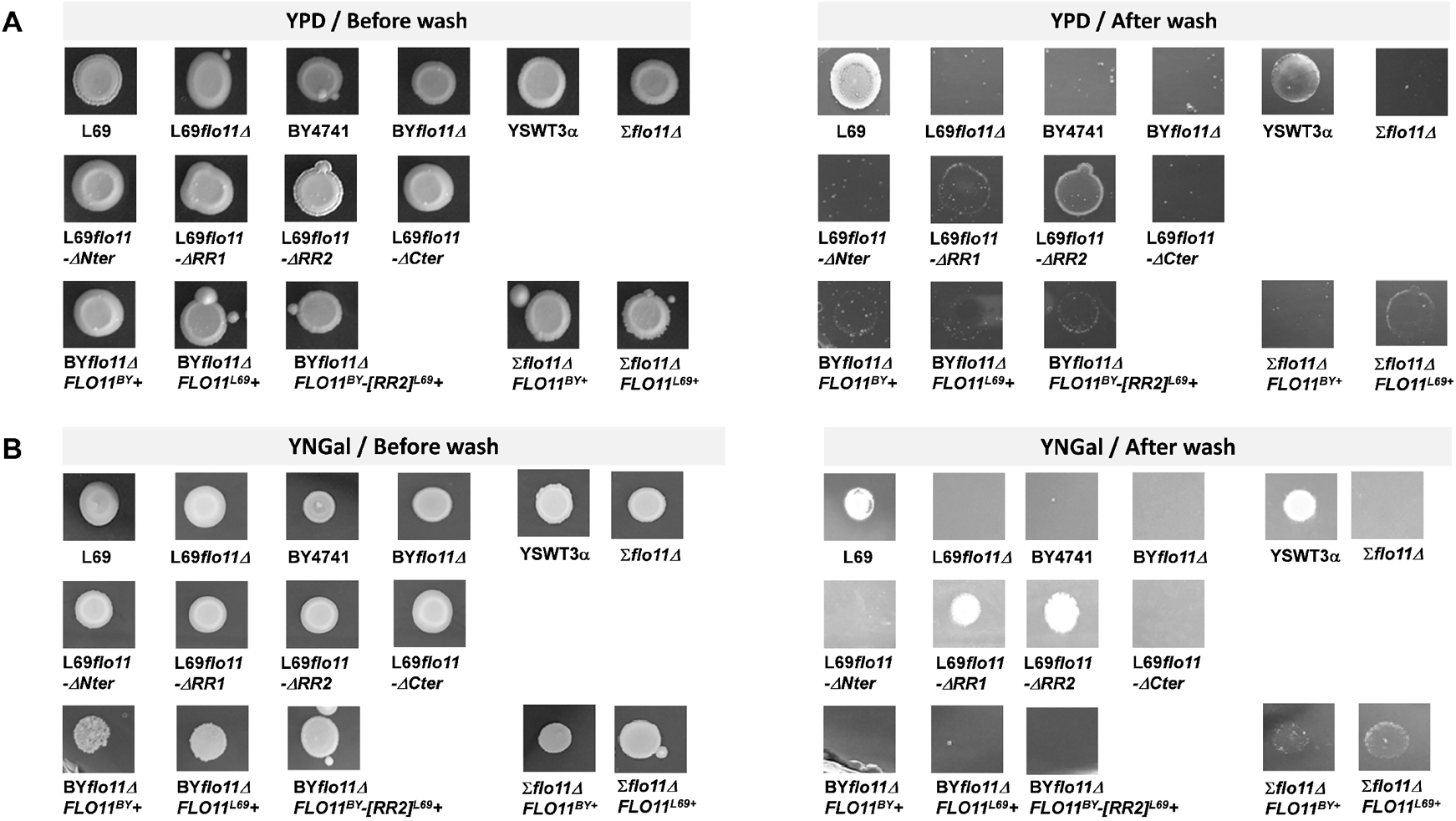
the Invasive growth in agar is abolished by deletion of either N or C-terminal of the Flo11p. All strains were pre-grown in YNGal that was supplemented with the auxotrophic requirements when needed (*ie* uracil, leucine, histidine, methionine at 0.1% for BY4741, BY*flo11Δ* and YSTW3α but only leucine, histidine, methionine for BY*flo11Δ* and Σ*flo11Δ* expressing *FLO11*^*BY*^, *FLO11*^*L69*^ or *FLO11*^*BY*^*-[RR2]*^*L69*^) until stationary phase. Then, 10 µL of these cultures were spotted on 2% (w/v) agar plate made with YPD (Panel A) or YNGal complemented with auxotrophic requirements (panel B). Plates were incubated at 30°C for 8 days and washed under a stream of water. They were photographed before and after washing.

We then investigated which domains or sequences of the Flo11p^L69^ are necessary for the *in vivo* invasive growth phenotype. Results in **Figure 9** show that the invasion in agar was completely lost in strains expressing a Flo11p that lacks either the N or the C-terminal domain. This phenotype was reduced in L69*flo11-ΔRR1* strain cells, whereas the removal of the RR2 sequence (L69flo11-ΔRR2) had barely no effect on the capacity of cells to invade agar (**Figure 9** and see also **Figure S12**). From these results, it can be concluded that RR2 region, which is needed for the formation of adhesive nanodomains, has no role in the invasion growth phenotype. More puzzling results were obtained with the laboratory strain BY4741 transformed with *FLO11*^*L69*^ or *FLO11*^*BY*^ carried on a 2µ plasmid under the *GAL1* promoter (pYES2.1). Indeed, we found that this strain failed to invade agar in spite of huge ectopic overexpression of *FLO11*^*BY*^ or *FLO11*^*L69*^ as in YNGal plates lacking uracil (YNGal Ura^-^) (**Figure 9B**) under which these genes were shown to be exceedingly transcribed (see Figure S2). On the other hand, the invasiveness of the haploid YSWT3α deleted for *FLO11* (Σflo11Δ) transformed with pYES2.1 plasmid carrying either *FLO11*^*BY*^or *FLO11*^*L69*^ was clearly discernable in the selective YNGal Ura^-^(**Figure 8B**), with *FLO11*^*L69*^ being slightly more effective than *FLO11*^*BY*^. Nonetheless, the invasive phenotype of Σflo11Δ τhat overexpressed *FLO11*^*BY*^ or *FLO11*^*L69*^ was weaker than that of this strain expressing its endogenous Flo11 protein.

### Flo11 protein and the cell wall integrity

Flo11p is a highly mannosylated cell surface protein that is retained to the cell wall inner network of β-1,6 glucans through a GPI anchor (Klis *et al*., 2006). We found that the loss of function of this gene resulted in a higher sensitivity to drugs such as Calcofluor White (CFW), caffeine and Congo Red (CR) that are commonly used to assess integrity of the yeast cell wall (Levin, 2005). Furthermore, we found that the higher sensitivity to these cell wall drugs was mainly associated with the lack of the C-terminal region (from aa 1297 to 1721, see **Fig. 3**) of Flo11p although removal of RR1 or RR2 sequence of Flo11^L69^ protein resulted in mutant strains that were also slightly more sensitive than wild type (see **Figure S13**). The higher sensitivity of L69*flo11-ΔCter* strain could be explained by the lack of retention of the protein at the cell surface due to the removal of the GPI anchor. Under glycosylation of Flo11p could be invoked for the slight increase of sensitivity to cell wall drugs of strains expressing variants of Flo11^69^p that have been deleted for RR1 since the repeated sequences in this region are thought to be heavily glycosylated and cell surface glycosylation is important in the cell wall integrity (Strahl-Bolsinger *et al*., 1999, Free, 2013).

## DISCUSSION

In this work, we have demonstrated that the abundant and dense patches observed by AFM on the cell surface of the industrial wine yeast L69 strain (Schiavone *et al*., 2015) correspond to adhesion nanodomains, with similar physical characteristics as those formed on the cell surface of the pathogen *C. albicans* (Alsteens *et al*., 2010, Formosa *et al*., 2015). We moreover showed that the flocculin encoded by *FLO11* is the sole protein involved in the formation of these nanodomains, and this event can be prevented upon treatment of the cells with anti-amyloid peptides or anti-amyloidophilic dyes. Altogether, these data are in line with the model of Lipke *et al*. (Lipke *et al*., 2012, Lipke *et al*., 2018) arguing that adhesion nanodomains require wall-anchored proteins that must harbour in their protein sequence the following features: (i) amyloid-β-aggregation prone sequences of 5 to 7 amino acid residues and (ii) serine/threonine-rich ‘T domain’ enriched of β-branched aliphatic amino acids Ile, Val and Thr. While all flocculins encoded by *FLO1, FLO5, FLO9* and *FLO11* in yeast display these criteria, it was intriguing to know why such dense and abundant nanodomains had not been observed so far in *Saccharomyces cerevisiae*, even though physical *FLO*-dependent-modification of cell surface upon hydrodynamic shear force have been recorded (Chan & Lipke, 2014, Chan *et al*., 2016). The data reported in this work clearly showed that Flo11 protein from L69 strain differs from that of *Saccharomyces* strains sequenced so far, including the laboratory strains BY4741, Σ1278b and the Flor strain 113d (Fidalgo *et al*., 2006), by having a unique 230 amino acids sequence termed RR2 near the C-terminus. Interestingly, this extra sequence is likely the result of a duplication of a region of about 115 amino acids located at the C-terminal of Flo11 flocculins, which is characterized by the presence of β-aggregation sequence “VVSTTV” followed by “ITTTFV”. This duplication thus provides 2 additional amyloid-core sequences, which are clearly essential in the process of nanodomains formation in response to a tensile force, as demonstrated by the insertion of RR2 sequence into the Flo11p of BY4741 enabling this strain to form adhesion nanodomains with similar physical properties as those formed on the cell surface of L69 strain.

The existence of this unique RR2 sequence in Flo11p prompted us to investigate the importance of amyloid forming sequences in the cell surface properties conferred by this protein, and by extension on the role of A, B and C domains in the *in vivo* function of Flo11p in the yeast *Saccharomyces cerevisiae*. Firstly, we found that adhesion nanodomains can be formed only in yeast cells expressing the whole Flo11 protein, which supports the notion that this protein must remain attached to the cell wall and must be unfolded through its tandem repeats upon the action of an extension force in order to expose amyloid-core sequences for interactions between neighbouring flocculin molecules, according to the model of Lipke *et al*. (Lipke *et al*., 2012). Secondly, we showed that each domain of Flo11p contributes to cell-cell aggregation and that this phenotype is potentiated by increasing the number of amyloid-forming sequences in Flo11p. The finding that this phenotype was not abrogated when domain A was deleted could be at first glance in contradiction with the data of Kraushaar *et al*. (Kraushaar *et al*., 2015) who showed that this domain is essential for conferring cell-cell adhesion by facilitating homotypic interactions. However, the lack of A domain can be in part overcome by amyloid-forming sequences as these later could promote clustering of Flo11 molecules in *cis*, which enables cell-cell adhesion by trans-interaction as proposed by these same authors (Kraushaar *et al*., 2015, Bruckner *et al*., 2020). As a consequence, one can predict a complete loss of cell-cell aggregation would require ablation of both A-domain and RR2 in Flo11p. However, in spite of strong cell-cell aggregation, L69 strain did not exhibit a flocculation phenotype (Verstrepen & Klis, 2006). However, this phenotype is likely more a curiosity displayed by *S. cerevisiae var. diastaticus* and not expressed by other yeast strains in spite of their high expression of *FLO11*. Barua *et al*. (Barua *et al*., 2016) attributed the capability of *S. diastaticus* to flocculate to a lack of a 15 amino acids sequence starting at residue 115 (NTDWIDNPLVSRCDE), the presence of which reduces homotypic interactions characteristic of flocculation. However, this explanation is likely incorrect because not solely this sequence does not exist in Flo11p of L69 and BY4741 strains (see **Figure S3** in supplementary data), but, in addition, Bruckner et al (Bruckner *et al*., 2020) showed that this 15 amino acids sequence actually enhanced cell-cell adhesion.

Cell surface hydrophobicity and adherence to polystyrene surface are two well-established properties that Flo11p confers to yeast cells. Here, we showed that these two phenotypes are mainly dependent on the abundance of Flo11p at the surface of the yeast cell, whatever their strain origin. Nevertheless, the fact that the intensity of plastic adhesion was dependent on pH was not unexpected, except that it was surprising to find that this dependence was not solely attributed to the N-terminal that carries A-domain of Flo11p, as reported by (Kraushaar *et al*., 2015). In our hands, we found no role of A-domain in adherence to plastic at pH 5.0, whereas this domain has a major role at pH 8.0. How can we reconcile these apparent discrepancies? A likely explanation would be to consider that the contribution of Flo11p to plastic adherence at pH 5.0 is relatively weak, whereas this contribution is proportionally higher at pH 8.0. Arguments in favour of this suggestion are found in the fact that adherence of BY4741 at pH 5.0 is independent to *FLO11*, that the drop in adherence upon loss of *FLO11* is more pronounced at pH8.0, and that adherence of YSWTα strain at pH 8.0 is much higher than at pH 5.0. On the other hand, Kraushaar et al (Kraushaar *et al*., 2015) could not make this observation because they were working with a strain in which *FLO11* gene was overexpressed, which clearly exacerbated adherence to plastic, and in consequence have masked effect of other cell wall proteins in this property. At variance to the role of N-ter in the adherence phenotype, the hydrophobicity property of Flo11p is dependent on a cooperation of various domains of this protein, although the N-terminus and the central B-domain contributed most to this property. These data were consistent with the fact that the N-terminal of Flo11p is highly hydrophobic, notably due to the presence of several tryptophan residues (Lo & Dranginis, 1998, Mortensen *et al*., 2007, Kraushaar *et al*., 2015) and that gain of hydrophobicity increases with the number of tandem repeat in the B-domain of Flo11p (Fidalgo *et al*., 2006, Zara *et al*., 2009). However, this wine strain did not elicit velum formation under a flor medium condition in spite of the fact that the number of repeats in Flo11p of L69 strain is similar to that of Flo11p from the flor yeast strain 133d (46 vs 49) and higher than Flo11p of S288c or Σ1278b (39 and 23, respectively). Two possible, and nonexclusive, possibilities that remain to verify could be invoked to account for this failure of L69 strain as well as other yeast strain to form velum. This can be due to the nature of the repeats as the FLO11 gene of strain 133d presents a single type of sequence repeat of 81 nt whereas all other *FLO11* harbour at least 4 different sequence repeats. Alternatively or complementary, Flo11p of strain 133d is more glycosylated than that of Flo11p of strain 69 in spite of the protein have similar number of repeats.

A dominant phenotype brought about by expression of *FLO11* is the ability of haploid yeast cells to undergo invasive growth in agar or pseudohyphal growth for a diploid cell (Gancedo, 2001). In this work, we found that L69 exhibited a very intense invasive growth phenotype, which was at first glance surprising since L69 strain is diploid. This phenotype is commonly not expressed in diploid cells unless *FLO11* is highly expressed (Lo & Dranginis, 1998), supporting our previous data that this gene is highly expressed in this strain (Schiavone *et al*., 2015). We moreover showed that the N-ter and the C-ter of Flo11p are indispensable to elicit agar invasion phenotype. While it was already reported that invasion in agar was abrogated upon deletion of the N-terminus that carries the A-domain of Flo11p (Kraushaar *et al*., 2015), the importance of C-terminus in this phenotype had never been reported. This result could be explained by the inability of the cell to remain trapped in the agar due to the fact that Flo11p, in the absence of its C-terminus, can no longer remain attached to the cell wall network. On the other hand, the invasive growth in agar was strongly diminished upon deletion of repeat sequences in the B-domain. This result can be explained by either one of these two possibilities or a combination of both. Either, it is due to a reduction of glycosylation of Flo11p as a defect in this process was found to reduce invasive growth in agar (Meem & Cullen, 2012). Alternatively, the removal of tandem repeat in B-domain (*ie* RR1 sequence) may prevent Flo11p from reaching the cell surface and therefore reduce the capacity of the cells to remain trapped in the agar. Finally, this study brought to light new data on the process of invasive growth. On the one hand, we found that invasion in agar of L69 strain was equally intense under any kind of growth media, whereas this phenotype in the well-established Σ1278b background strain was weak in a galactose agar medium and absent in a glucose agar synthetic medium. In addition, the efficient agar invasion phenotype in L69 strain could not be solely attributed to *FLO11* gene because the overexpression of this gene in YSTW3a *flo11Δ* (derivative of Σ1278b) did not provide to this strain a higher capacity to invade agar than that of the isogenic YSTW3α expressing its endogenous gene. On the other hand, the ectopic overexpression of *FLO11* in BY4741 did not rescue the inability of this haploid strain to elicit invasive growth phenotype. Altogether, these data indicated that the Flo11p-dependent invasive phenotype involves additional genetic factors that are absent in BY4741 and that are more expressed in L69 strain, accounting for the higher potency of this strain to invade agar.

## MATERIALS AND METHODS

### Strains and growth conditions

The *Saccharomyces cerevisiae* strains used in this work are listed in Table S4. Unless otherwise stated, strains were cultivated in rich YPD medium at 30°C (1% w/v yeast extract, 1% w/v bactopeptone and 2% w/v glucose), When using synthetic YNGlu or YNGal media (0.17% w/v yeast nitrogen base, 0.5% w/v (NH_4_)_2_SO_4_, with 2% w/v glucose or 2% w/v galactose), they were supplemented with appropriate amino acids at 0.1 % (w:v). YNB Acetamide medium (0.17% w/v yeast nitrogen base, 0.66% w/v K_2_SO_4_, 0.06% w/v acetamide and 2% w/v glucose) was used for the selection and propagation of L69 mutants transformed with plasmid bearing amdSYM cassette (Solis-Escalante *et al*., 2013) used for CRISPR-Cas9 deletion (Ryan *et al*., 2016). YNB Fluoroacetamide medium (0.17% w/v of yeast nitrogen base, 0.5% w/v of (NH_4_)2SO_4_, 0.23% w/v fluoroacetamide and 2% w/v glucose) was used to excise the amdSYM cassette from the genome of L69 generated mutants. For solid media, agar was added at 2% (w/v) before sterilization at 120°C for 20 min.

### Plasmids and strains construction

To overexpress *FLO11* gene or any of its alleles in yeast strain, expression vector pYES2.1 TOPO TA (Table S5) was used which carries *GAL1* as promoter and *CYC1* terminator. The *FLO11* ORF from BY4741 (*FLO11*^BY^) and L69 (*FLO11*^L69^) were amplified by PCR using the primers FLO11_TOPO_f and FLO11_TOPO_r (Table S6) and ligated into pYES2.1 TOPO TA vector (ThermoFisher Scientific) between *GAL1* promoter and *CYC1* terminator according to the manufacture’s protocol. Correct integration was confirmed by restriction digestion and plasmid sequencing.

Deletion of *FLO11* were constructed in L69 strain using CRISPR-Cas9 strategy (Ryan *et al*., 2016). The high copy pCas9-amdSYM plasmid (derived from pML107, Addgene) that constitutively expresses the gene encoding Cas9 endonuclease protein and carrying a gRNA expression cassette (Sap1 cloning sites) was used. It confers the ability to *S. cerevisiae* to use acetamide as the sole nitrogen source (Solis-Escalante *et al*., 2013) enabling selection of prototroph strains. As the CRISPR/Cas9 technique requires the identification of a unique 20N (NGG) sequence, the candidate target sequence was identified using the online software CRISPR-direct (https://crispr.dbcls.jp), and determined as to be located in the middle of each region of *FLO11* to be deleted. Linear healing fragments of 120 bp were designed with 60 bp overlapping the upstream and downstream sequences of the region to be deleted, namely the N-terminal (from 4 bp to 675 bp), the C-terminal (from 3889bp to 5163 bp), the RR1 domain (from 735bp to 2573bp) and the RR2 domain (from 3575bp to 4375bp). Ligation of gRNA sequence into the plasmid was made using a T4 DNA ligase at 16°C overnight and transformed into DH5α chemically competent bacteria according to the manufacturer protocol (NEB). The presence of the new gRNA into pCas9-amdSYM was confirmed by Sanger sequencing using the M13 forward primer. The generated plasmids were used to transform L69 strain and healing fragments were added to repair the double strand break made by CRISPR-Cas9 protein according to the wanted deletion. Constructions were verified by PCR amplification and Sanger sequencing (Supplementary data, Table S6).

For BY*flo11Δ FLO11*^*BY*^*-[RR2]* ^**+**^ strain, a chimeric gene constituted of the N-terminal and central regions of *FLO11* from BY4741 strain fused to the C-terminal domain of *FLO11* from L69 (including the RR2 region) was constructed. To this end, a 2967 bp fragment was PCR amplified from *FLO11*^BY^ using the primers FLO11_TOPO_f and FLO11_BY_2964_r (see Table S3), whereas a 1794 bp fragment was PCR amplified from *FLO11*^*L69*^ using the primers R269 inFLO11BY_f and FLO11_TOPO_r (Table S3). Both fragments share a 30 nucleotides sequence overlapping that allow their fusion by overlapping PCR reaction. The resulting 4731 nucleotides fragment was PCR amplified using FLO11_TOPO_f and FLO11_TOPO_r oligonucleotides (Supplementary data, Table S6). The resulting chimeric gene *FLO11*^*BY*^*-[RR2]*^*L69*^ was ligated into pYES2.1 TOPO TA vector as described above.

Yeast strains were then transformed with each of the resulting plasmids using yeast transformation procedure as described in (Gietz & Schiestl, 2007)

## Bioinformatics

The gene *FLO11* was retrieved from the L69 strain genome which has been fully sequenced internally (Lallemand, Inc, unpublished data) using the Pacific Biosciences method, whereas the sequences of *FLO11* in the laboratory strain BY4741 and Σ1278b were uploaded from the SGD database (https://www.yeastgenome.org). Clustal Omega was used to perform amino acid sequence alignments (Sievers & Higgins, 2014). Structural and functional comparison between the sequences of Flo11p^L69^, Flo11p^BY4741^ and Flo11p_Σ_^1278b^ were carried out using Hydrophobic Cluster Analysis (HCA) (Lemesle-Varloot *et al*., 1990) giving a plot of each open reading frame that draws as a helical projection, vertically repeated. Secretion signal and the GPI signal anchorage to cell wall β-glucan were searched using SignalP-4.1 server (http://www.cbs.dtu.dk/services/SignalP-4.1/) (Nielsen, 2017), PredGPI tool (http://gpcr.biocomp.unibo.it/predgpi/) (Pierleoni *et al*., 2008). TANGO software (http://tango.crg.es/) (Fernandez-Escamilla *et al*., 2004) with default settings for pH, ionic strength, and temperature was used to determine Flo11p regions with β-aggregation potential superior to 30%. Intragenic repeats in the *FLO11* ORF were screened using the EMBOSS ETANDEM software (http://emboss.bioinformatics.nl/cgi-bin/emboss/etandem) (Rice *et al*., 2000). Criteria were selected those with a repetition length >10, a score >20, and a repetition conservation >60%.

### Atomic Force Microscopy

Cells were collected from exponential growth, washed twice in acetate buffer (18 mM CH_3_COONa, 1 mM CaCl_2_ and, 1 mM MnCl_2_, pH 5.2), and immobilized on polydimethylsiloxane (PDMS) stamps as described before in (Formosa *et al*., 2015). AFM experiments were recorded at room temperature using a Nanowizard III system (JPK-Bruker) and MLCT cantilevers (Bruker). The spring constants of each probe were systematically measured by the thermal noise method according to (Hutter & Bechhoefer, 1993) and were found to be in the range of 0.01-0.02 N.m^-1^. AFM height and adhesion images were recorded in Quantitative Imaging™ mode (JPK-Bruker) (Chopinet *et al*., 2013), and the maximal force applied to the cell was limited to 1 nN. For each condition, 3 independent experiments were performed and at least 12 cells were imaged. All results were analysed with the JPK Data Processing software. The adhesion values measured on cells were determined from the retract force-distance curves. The stiffness values (kcell) were determined as the slope of the linear portion of the force versus indentation curves according to: 

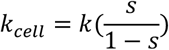

The effective spring constant of the cell *k* _*cell*_ is calculated from the experimental slope *s* of the force curve and the spring constant k of the cantilever measured by the thermal noise method. All stiffness and adhesion values were considered for the histograms, which were generated using OriginPro version 2020 (OriginLab, Northampton, USA). A synthetic peptide based on the amyloid sequence VVSTTV bearing a replacement of V by A (VASTTV) was purchased from Gencust and used as anti-amyloid peptide against Flo11p amyloid sequence. Cells were incubated for 90 minutes with 2 mg/ml of this peptide prior to AFM analysis.

### Quantitative Reverse Transcription PCR (RT-QPCR)

Unless otherwise stated, yeast strains were cultured in YPD medium and 5 DO of exponentially growing cells ((O.D._600nm_ ∼ 1.5) were harvested by centrifugation, nitrogen frozen and stored at −80°C. RNA extraction was carried out using RNEasy Plus Mini kit (Qiagen) according to manufacturer’s protocol. Quality and quantification of RNAs were determined using a Nanodrop 2000 (Thermo) and Bioanalyzer 2100 (Agilent). To synthetize cDNA, 1 µg of RNA was used in a 20 μL reaction mixture using the iScript cDNA synthesis kit (Bio-Rad). Reactions for RT-qPCR were performed in 20 µl reaction with 1 µl cDNA (0.25 ng.µl^-1^ final concentration), 10 µl of iQ SYBR Green Supermix buffer (Bio-Rad), 5 µl of Nuclease free water and 4 µl of the appropriate oligonucleotides (Table S2) at a final concentration of 0.25µM, designed using the qPCR Primer & Probe Design Webtool (https://www.eurofinsgenomics.eu/en/ecom/tools/qpcr-assay-design/), and run on a MyIQ real-time PCR system (Bio-Rad). All reactions were run in triplicate, with *TAF10* and *UBC6* used as reference genes (Teste *et al*., 2009). The relative transcript abundances of *FLO11* normalized to TAF10 and UBC6 were calculated based on the 2^-ΔΔCt^ method (Livak & Schmittgen, 2001).

### Assay of adherence to polystyrene

For determination of the adherence to polystyrene surfaces, the cells were grown overnight at 30°C in YN Gal (supplemented with auxotrophic requirements when necessary), washed once in PBS buffer and resuspended in YN Gal adjusted at either pH 5.0 or pH 8.0 with HCL or NaoH 1M to obtain 1 unit OD_600_. Aliquots of 100 μL of the culture were transferred into 96-well polystyrene plates and the cell suspensions were incubated at 30°C for 1 h at 200 rpm. An equal volume of 0.1% (w/v) crystal violet was then added to each well. After 15 min, the wells were washed 4 times with sterile water, and the adherence of the cells was quantified by solubilizing the retained crystal violet in 100 μL 95% ethanol. After 10 min the absorbance of the samples against the blank was measured at 595 nm in microplate reader (Biotek).

### Octane adhesion test

Hydrophobic features of yeast surface were determined by measuring their affinity for a nonpolar solvent as described in (Purevdorj-Gage *et al*., 2007). Overnight cultures were centrifuged at 2.000 *g* for 5 min and resuspended in fresh YN Galactose medium at OD_600_ of 1. After 3 h of static incubation at room temperature, O.D._600nm_ was measured (*A0*) and 1.2 ml of each culture was overlaid with 0.6 ml of octane (Sigma-Aldrich) in 15 × 100 mm borosilicate glass tubes. The tubes were vigorously vortexed for 2 min. and left on the bench for at least 15 min until complete separation of the two phases. A sample of the aqueous phase was taken with a Pasteur pipette and the O.D._600nm_ was measured (*A*). The results were expressed as the Octane adhesion index (% hydrophobicity), which represents the percentage of cells retained by the organic fraction, according to the equation: 

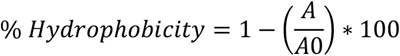

### Invasive agar growth assay

Agar invasion assays were performed according to the method previously described (Roberts & Fink, 1994). Strains were patched on glucose or galactose rich or synthetic medium plates supplemented with auxotrophic compounds when needed and grown at 30°C for 5 to 8 days. They were photographed before and after washing with distilled water.

### Cell-cell aggregation assay

Cells were grown in YNGal medium and harvested during entry in stationary phase. Cell concentration of each sample was adjusted to 8.10^7^ cells.mL^-1^ and 4μl were dropped on a microscope slide and observed with a microscope Eclipse 400 (Nikon) after being briefly vortexed at low agitation. Aggregates with at least 5 cells were counted and reported as a percentage as previously described in (Purevdorj-Gage *et al*., 2007).

### Other phenotypic assays

Sensitivity of yeast strains to caffeine (Sigma-Aldrich), calcofluor white (CFW; US Bio) and Congo Red (CR) was performed on YPD agar plates. Briefly, exponentially growing cells on YPD (O.D_600nm_ around 1) were collected by centrifugation and resuspended in sterilize water at 10 0D units. Series of 10-fold dilutions were spotted on YPD agar plates in the absence or presence of various concentrations of CFW or CR. Pictures were taken after 2 days of growth at 30 °C. Flocculation tests were carried out according to (Lo & Dranginis, 1996), starting with overnight yeast cultures in YPD, washed once with deflocculating medium (20 mM citrate pH 3.0 containing 5 mM EDTA), and resuspended in 1 ml of the same solution at 1.0 unit OD_600_. Then, CaCl2 (1M solution) was added at a final concentration of 20 mM and decrease of absorbance was monitored at 600 nm. Velum formation was carried out exactly as described in Zara et al. (Zara *et al*., 2005). The flor yeast strain A9 (kind gift from M. Budroni, Univ Sassari, Italy) was used as a control.

### Statistical analysis

All phenotypic assays were carried out at least 3 times with independent biological samples. Statistical analysis were made by one-way analysis of variance followed by Tukey’s test on Microsoft Excel software. Statistical significant values were denoted by asterisks on the figures as *= p-value < 0.05, ** =p-value < 0.01 and ***= p-value < 0.001.

## ACKNOWLEDGEMENTS

We are grateful Dr Jean Luc Parrou for advice on performing RT-qPCR experiments, to Dr Charlie Boone of University Toronto Canada, and Dr Marilena Budroni from University of Sassari, Italy, for the kind gift of yeast strains, and to Dr Mathieu Castex from Lallemand Inc. for continuous support on this work. CB is financed by ANRT (Agence Nationale de la Recherche et des Technologies) grant to carry out her PhD thesis.

## Supplementary data

**Figure S1:**
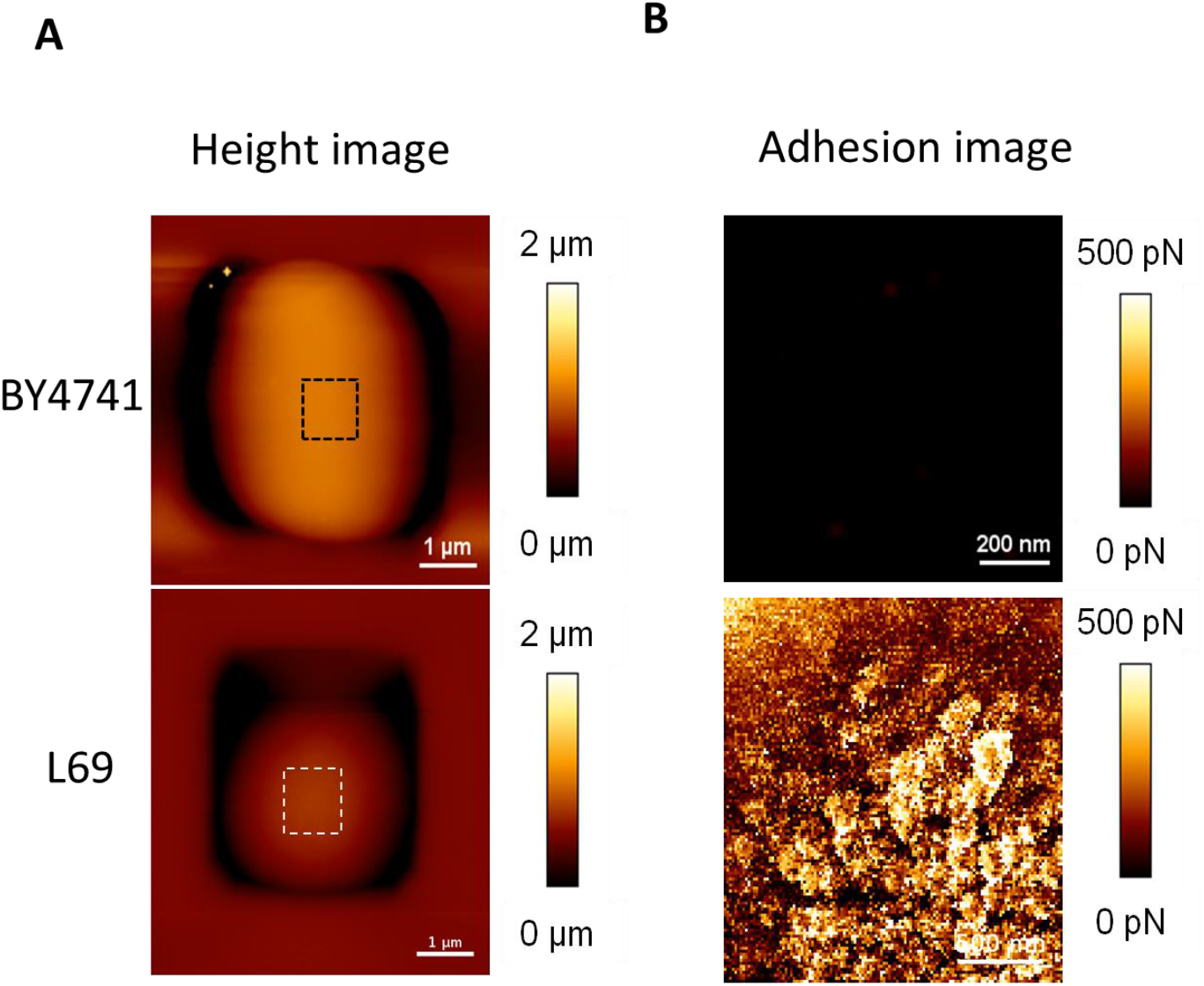
Cell surface analysis of BY4741 and L69 strain using silicon-nitride (Si3N4) AFM cantilevers. A single cell from strain BY4741 and L69 was trapped in the PDMS chamber as described in Material and Methods. AFM height image (z-scale: 2μm) (A) and adhesion image (z-scale: 500 pN) analyzed on the hatched square of the height image (B) were recorded in QI mode.

**Figure S2:**
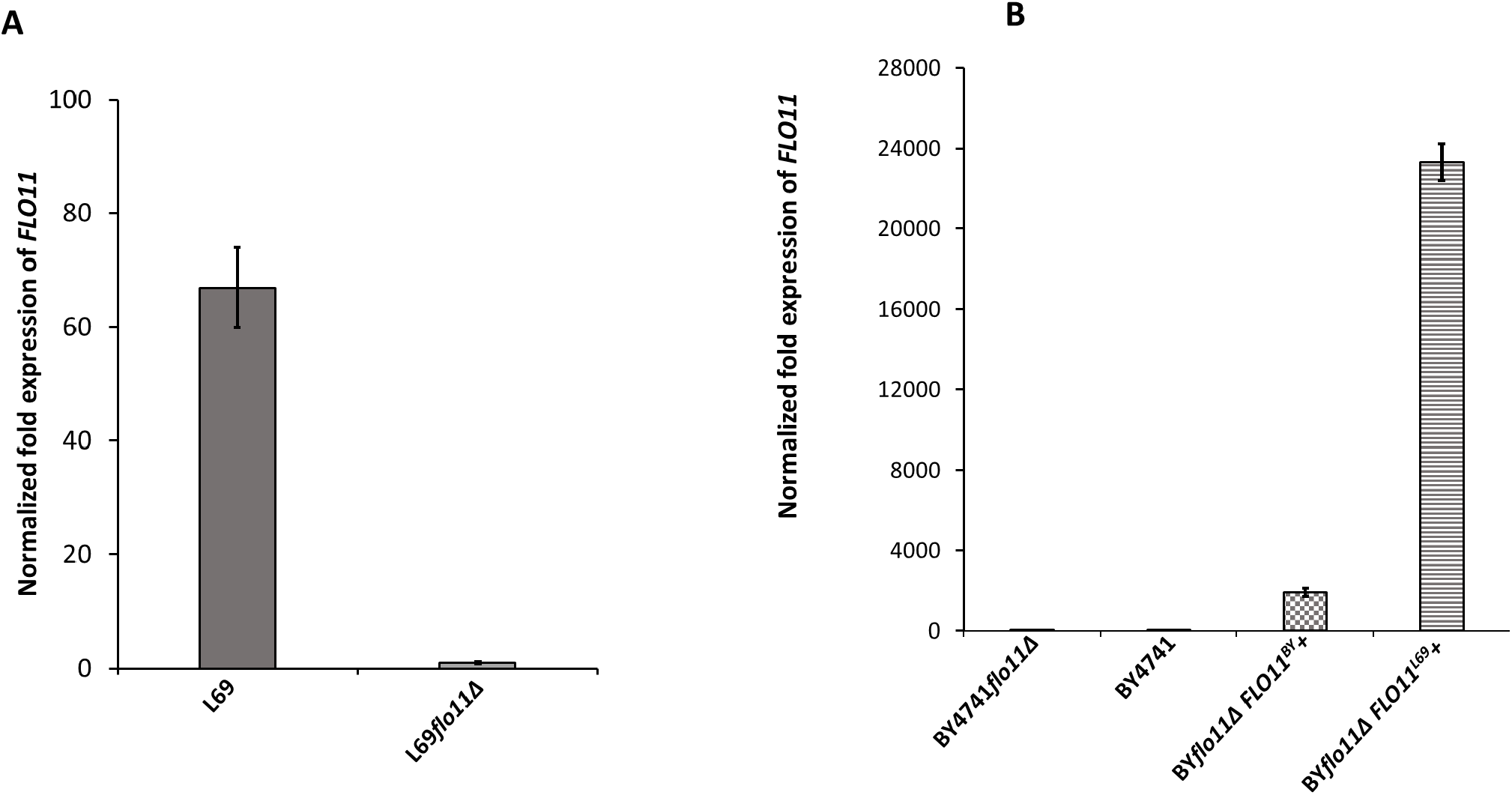
Expression levels of *FLO11* in L69 and BY4741 strain measured by RT-qPCR. Transcript levels of *FLO11* were determined in exponential cultures of L69 and BY4741 strains on YPD (A) or on YNGal supplemented with required amino acids, uracil and adenine at 0.1% (w/v), except for BY4741 transformed with *FLO11*^*BY*^ or *FLO11*^*L69*^ for which uracil was omitted. The transcripts was normalized to internal reference genes *TAF10* and *UBC6*.

**Figure S3:**
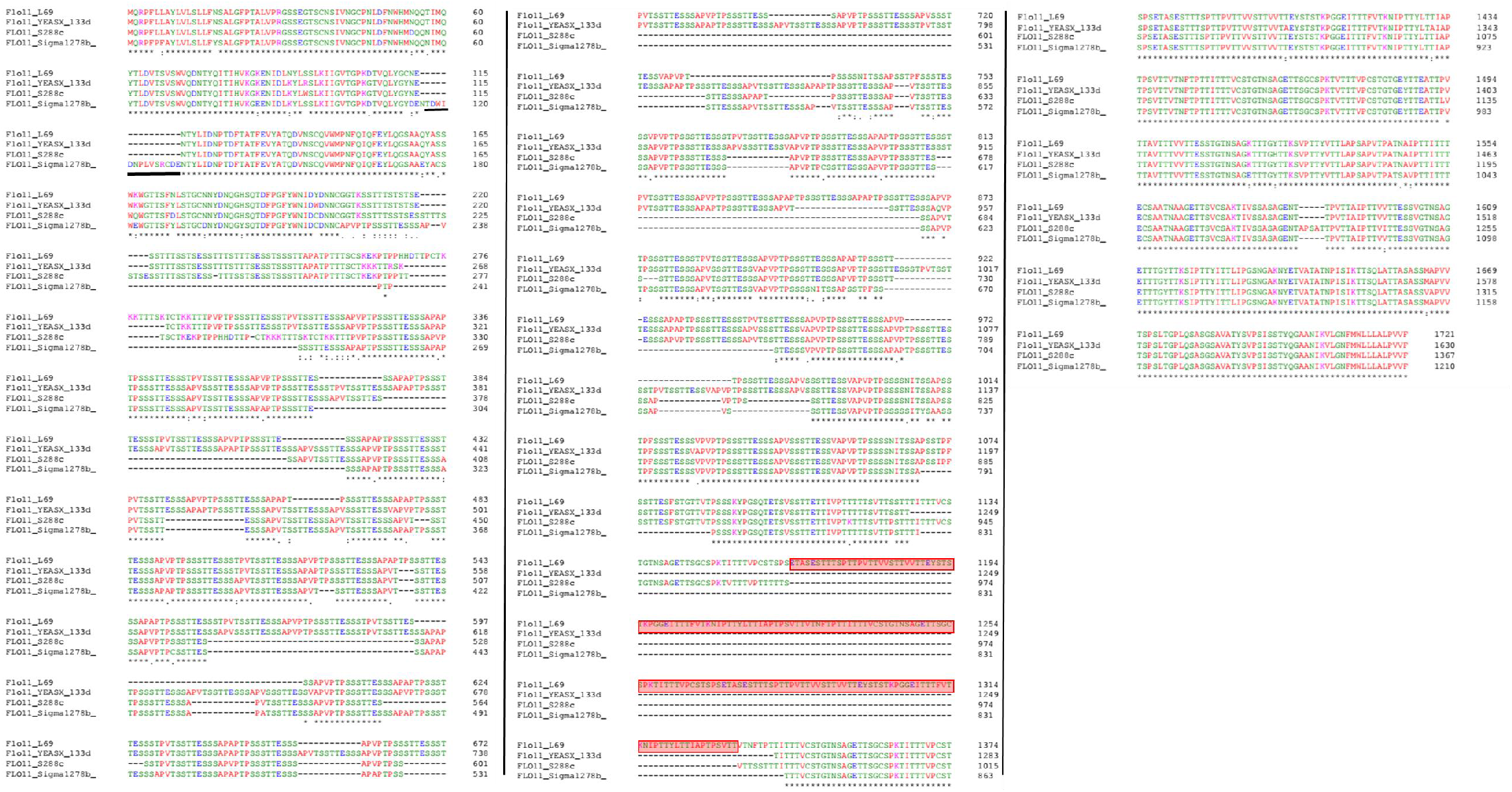
Sequences alignment of Flo11p from L69, BY4741, Σ1278b and 133d strains. The Flo11p sequences were aligned with Clustal Omega. The sequence of Flo11p from L69 strain was retrieved from the genome sequence of this strain (Lallemand Inc, unpublished data), whereas Flo11p from BY4741, Σ1278B and 133d were retrieved from public repository data at NCBI. Boxed in red indicate amino acid sequence (RR2) present in Flo11p of L69 strain and absent in Flo11p of the other strains. Asterisks indicate positions with identical amino acids, dots indicate single amino acid difference between amino acid sequences. Black bar highlights the unique 15 amino acid sequence in the Flo11 of Σ1278b strain

**Figure S4:**
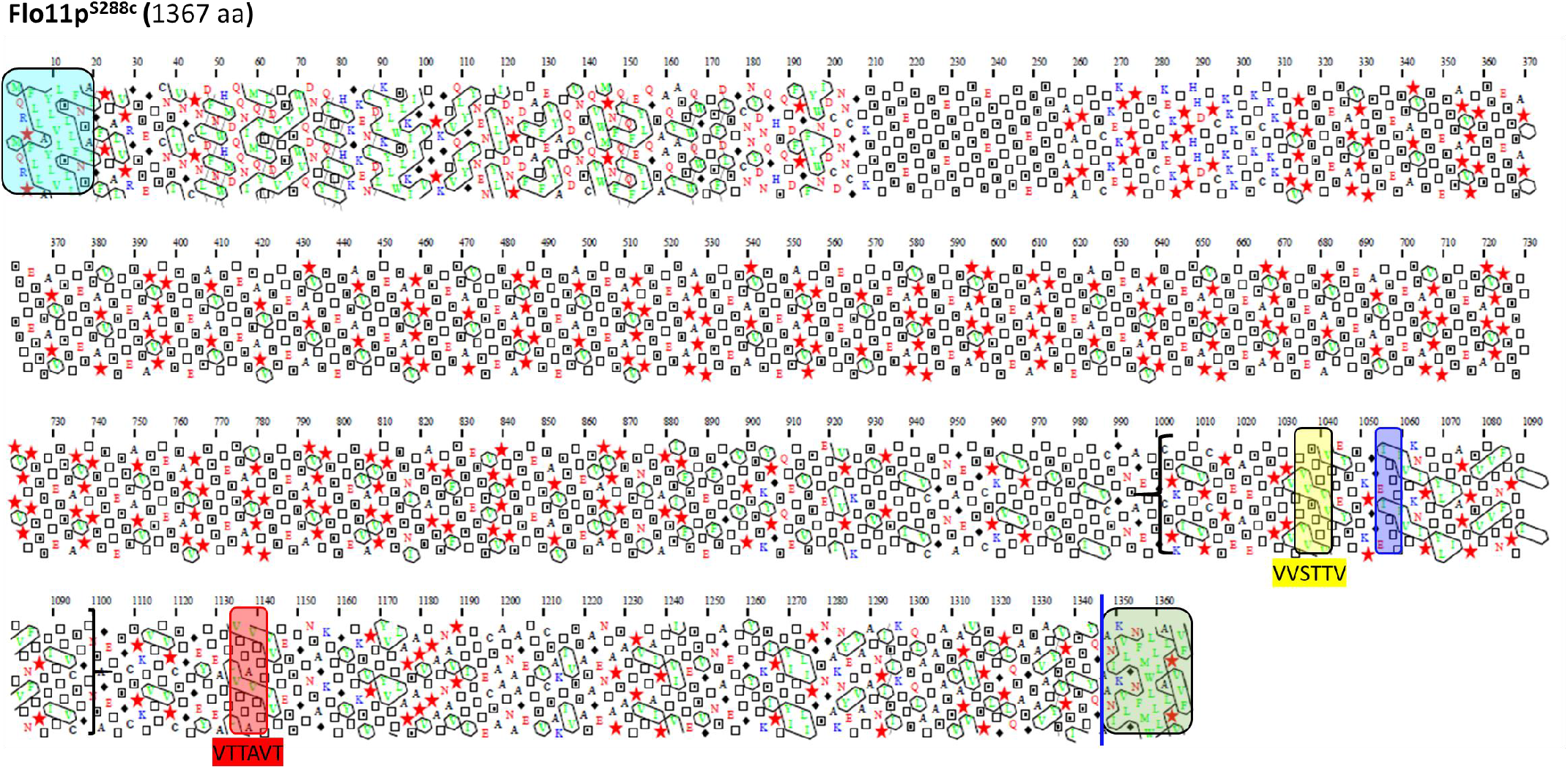
Schematic representation of the Flo11 protein sequence from BY4741 strain by Hydrophobic Cluster Analysis. The Flo11p from BY4741 strain is shown with the N-terminal secretion signal sequences boxed in blue determined with SignalP-4.1 server and the C-terminal GPI addition signals using PredGPI tool, boxed in green with a blue line indicating the omega-site position (GPI signal anchorage to cell wall β-glucan) which includes a β-aggregation motif FMWLLA. Amyloid core sequences ‘VVSTTV’ or ‘VTTAVT’ are highlighted in yellow and red boxes. The ITTTFV β-aggregation motif that specifically follows the first amyloid forming sequence is shown as violet box. Between bracket is the amino acids sequence that is found times repeated in Flo11p of L69 strain.

**Figure S5:**
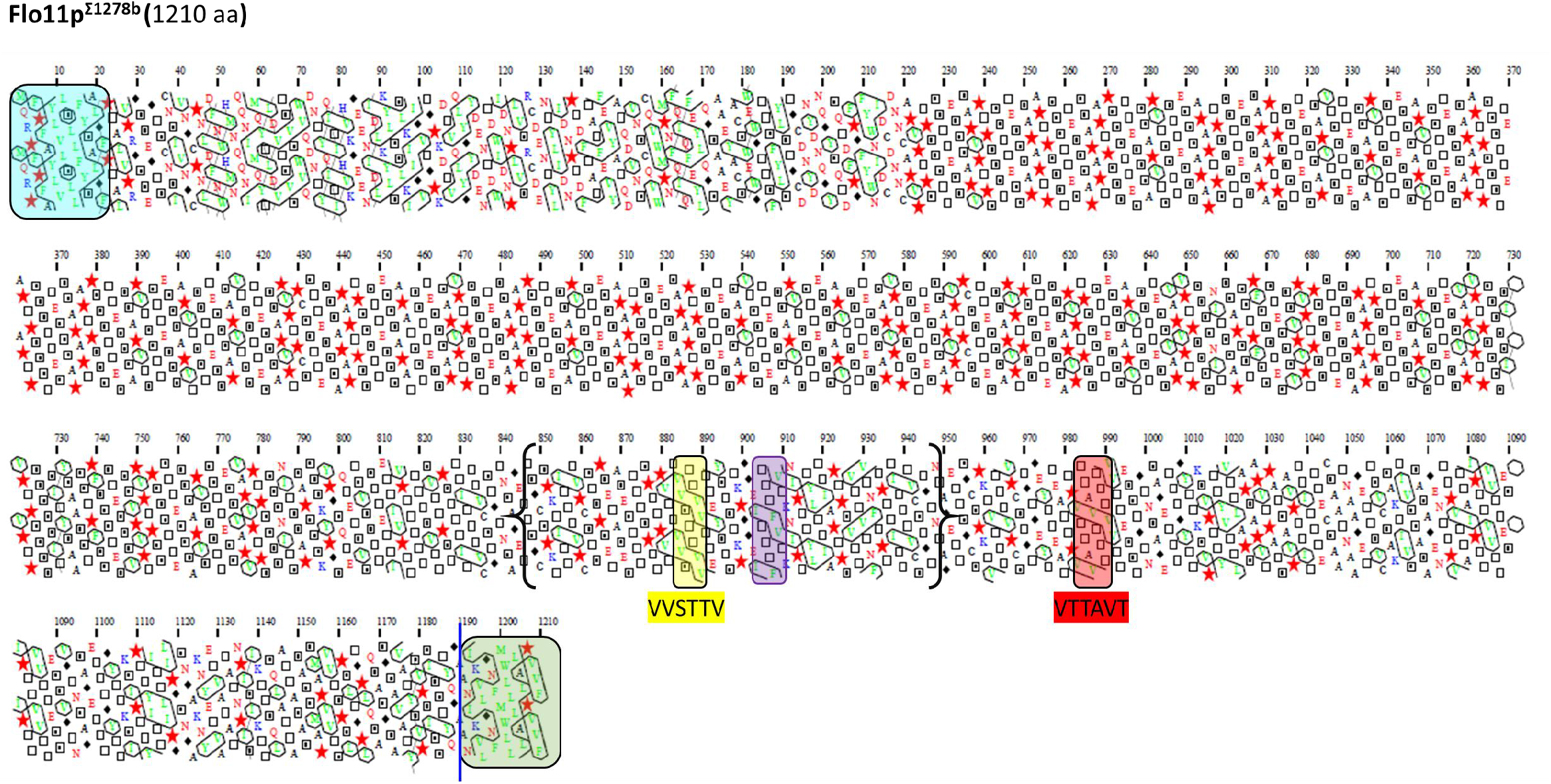
Schematic representation of the Flo11 protein sequence from Σ1278b strain by hydrophobic Cluster Analysis. Same description as in Fig.S4

**Figure S6:**
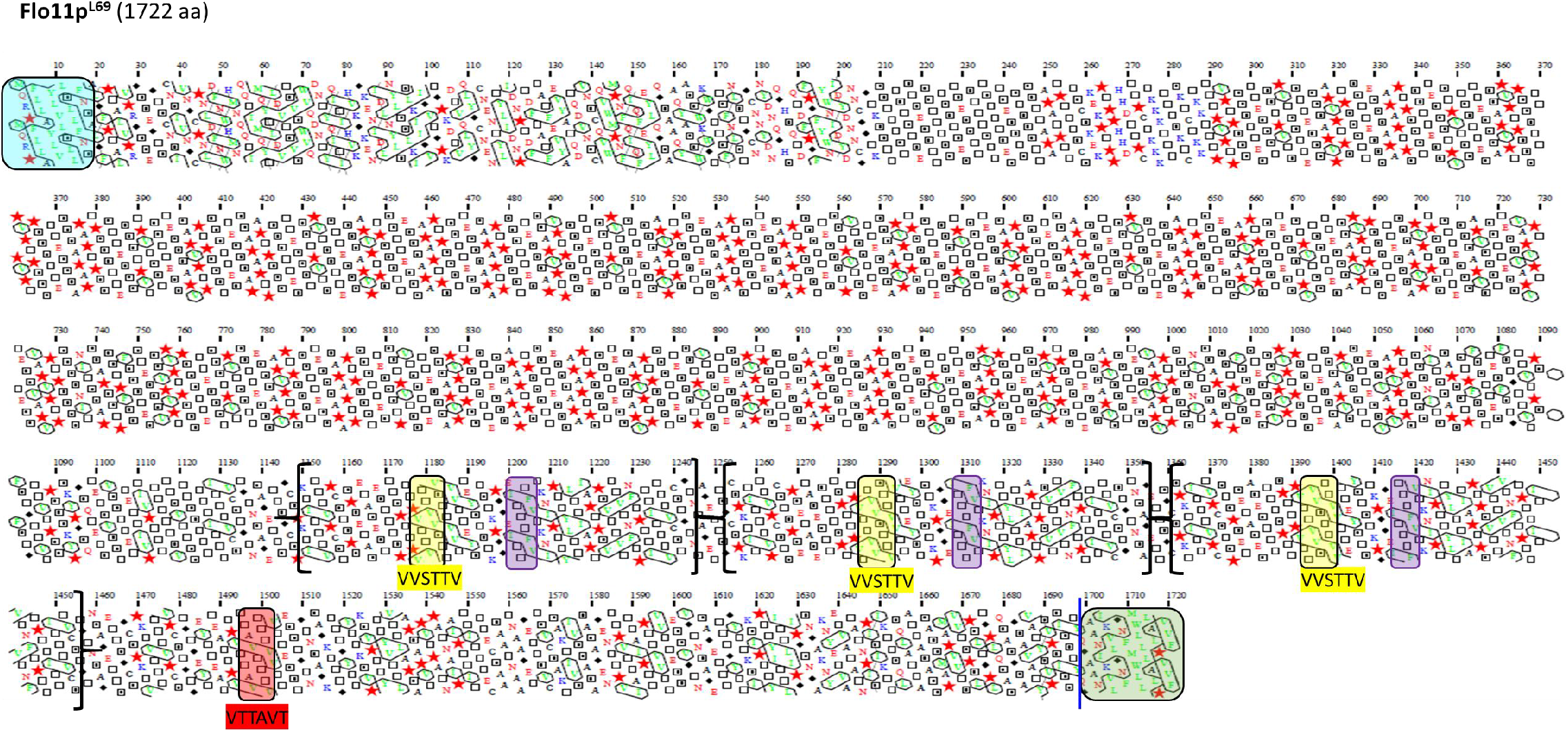
Schematic representation of the Flo11 protein sequence from L69 strain by Hydrophobic Cluster Analysis (HCA). Same as in Figure S4. Notice the presence of two additional amyloid-forming sequences, together with two additional β-aggregation motifs ITTTFV in this protein that are not present in Flo11p of all the other strains. In brackets is indicated the region that has been duplicated in the Flo11p to yield these two additional amyloid β-aggregation prone sequences in the Flo11p of L69 strain.

**Figure S7:**
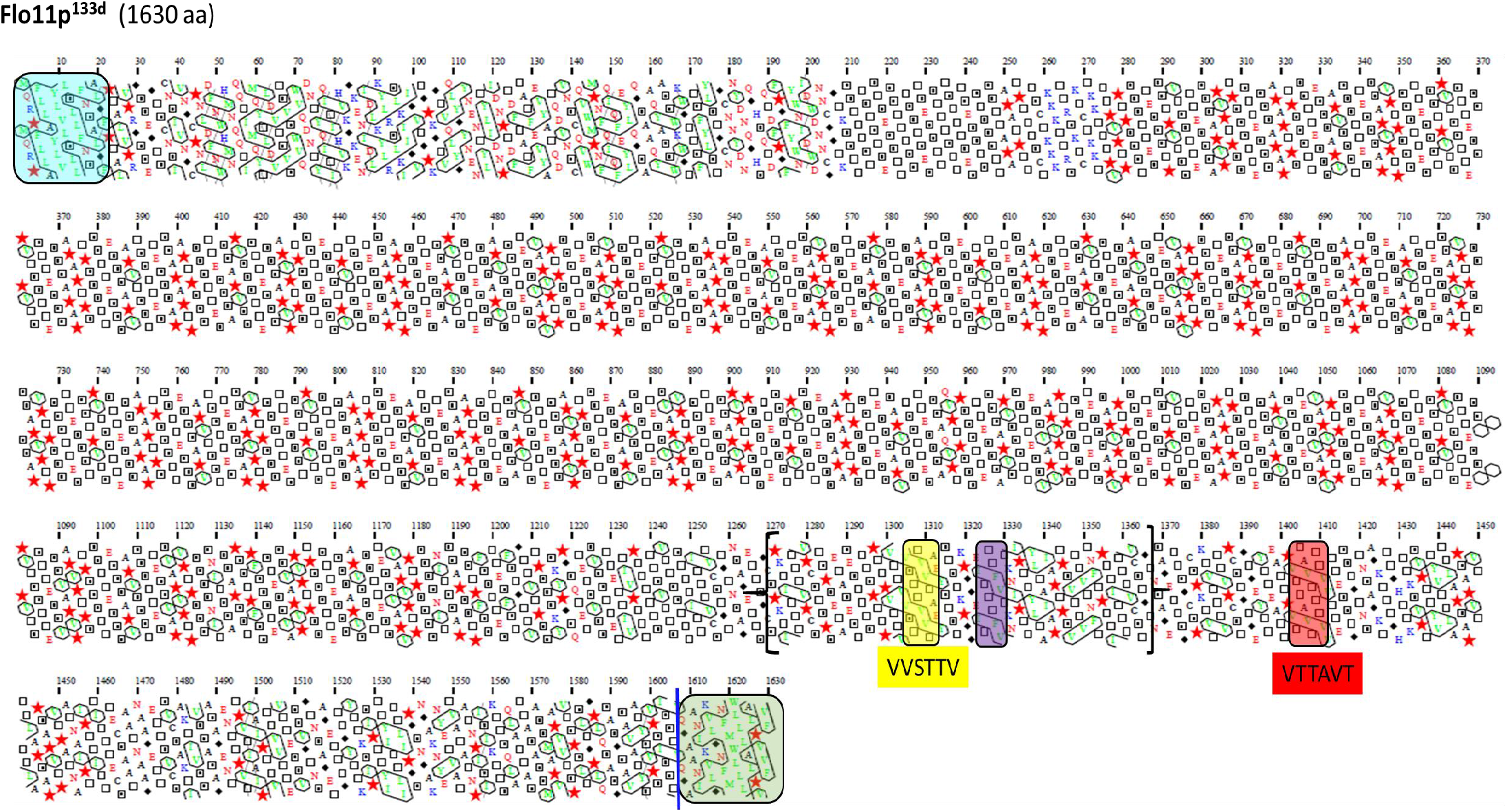
Schematic representation of the Flo11 protein sequence from flor yeast strain 133d by Hydrophobic Cluster Analysis. Same as in Fig.4

**Figure S8:**
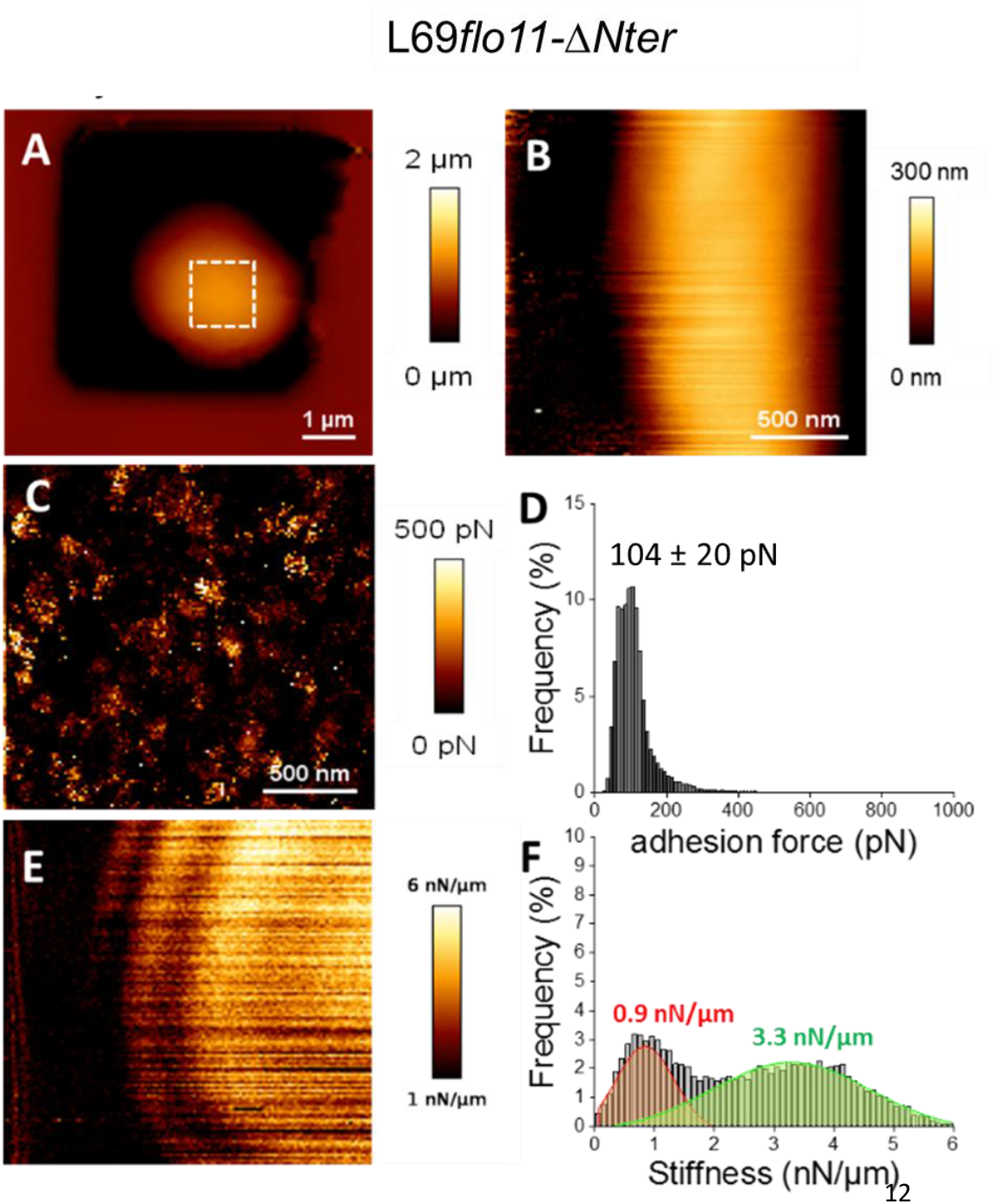
Cell surface analysis of the L69*flo11-ΔNter* strain using silicon-nitride (Si3N4) AFM cantilevers. AFM height image (A, B), adhesion image from the hatched square in A (C) and stiffness image (E) of a single yeast cell from L69 flo11-ΔNter trapped in a PDMS chamber is shown. In (D) is reported the adhesion force histogram obtained from 1024 force-distance curve recorded in QI™ mode on the area of cell surface illustrated in (C). In (F) is reported the stiffness histogram from 1024 force-distance curve recorded in QI™ mode. See Fig 2 and Material & Method for the visualization and quantitative determination of adhesion force and stiffness from the force-distance curves.

**Figure S9:**
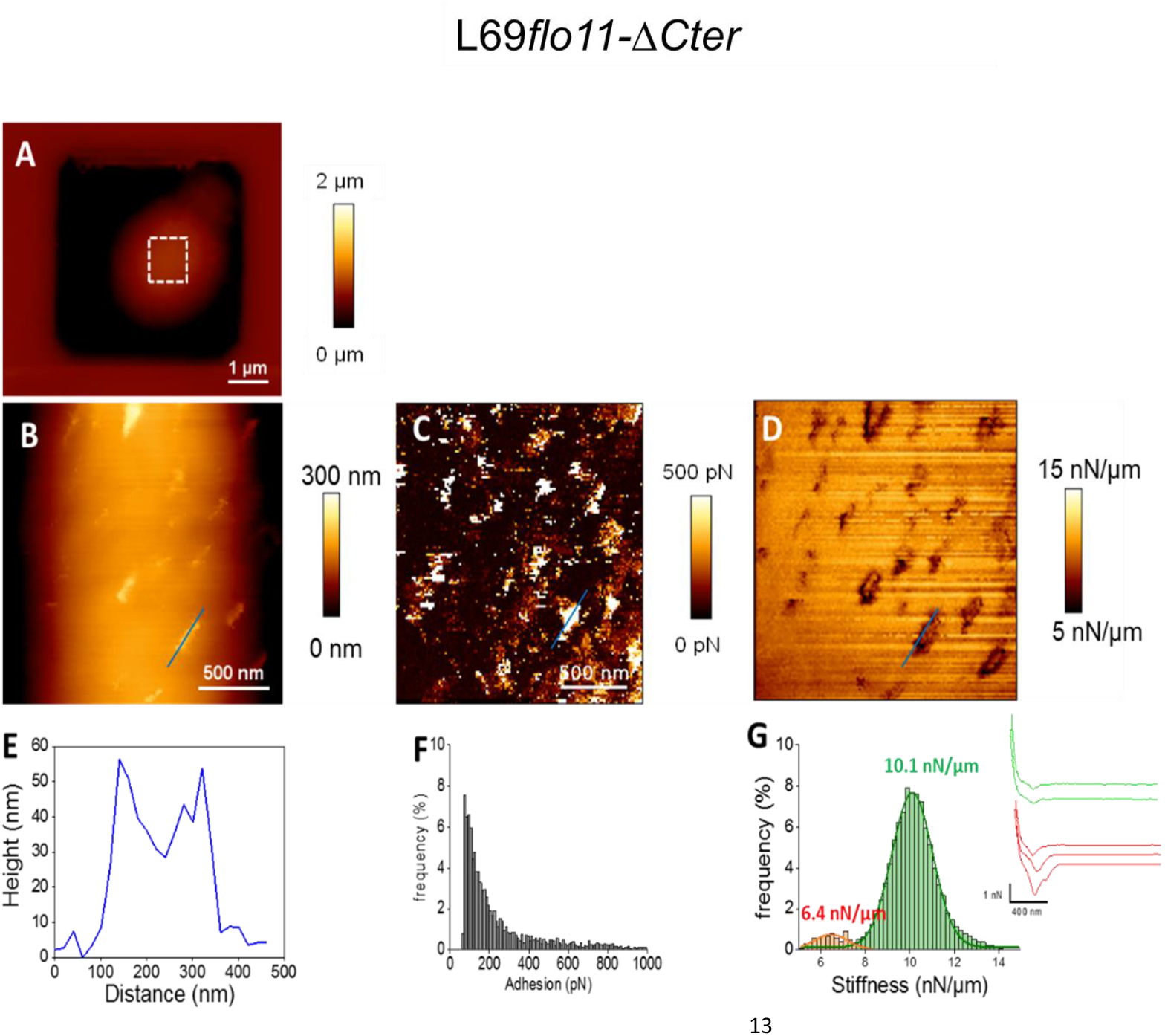
Cell surface analysis of the L69*fl011*-*ΔCter* strain using silicon-nitride (Si3N4) AFM cantilevers. AFM height image (A, B), adhesion image (C) and stiffness image (D) of a single yeast cell from L69 flo11-ΔCter is shown. In (E) is shown the height of the patches versus their size as indicated by blue line in the adhesion image in (B). in (F) is reported the adhesion forces histogram obtained from 1024 force-distance curve recorded in QI™ mode and in (G) the corresponding stiffness, with illustration of a few distance curves on the left of the (G) figure.

**Figure S10:**
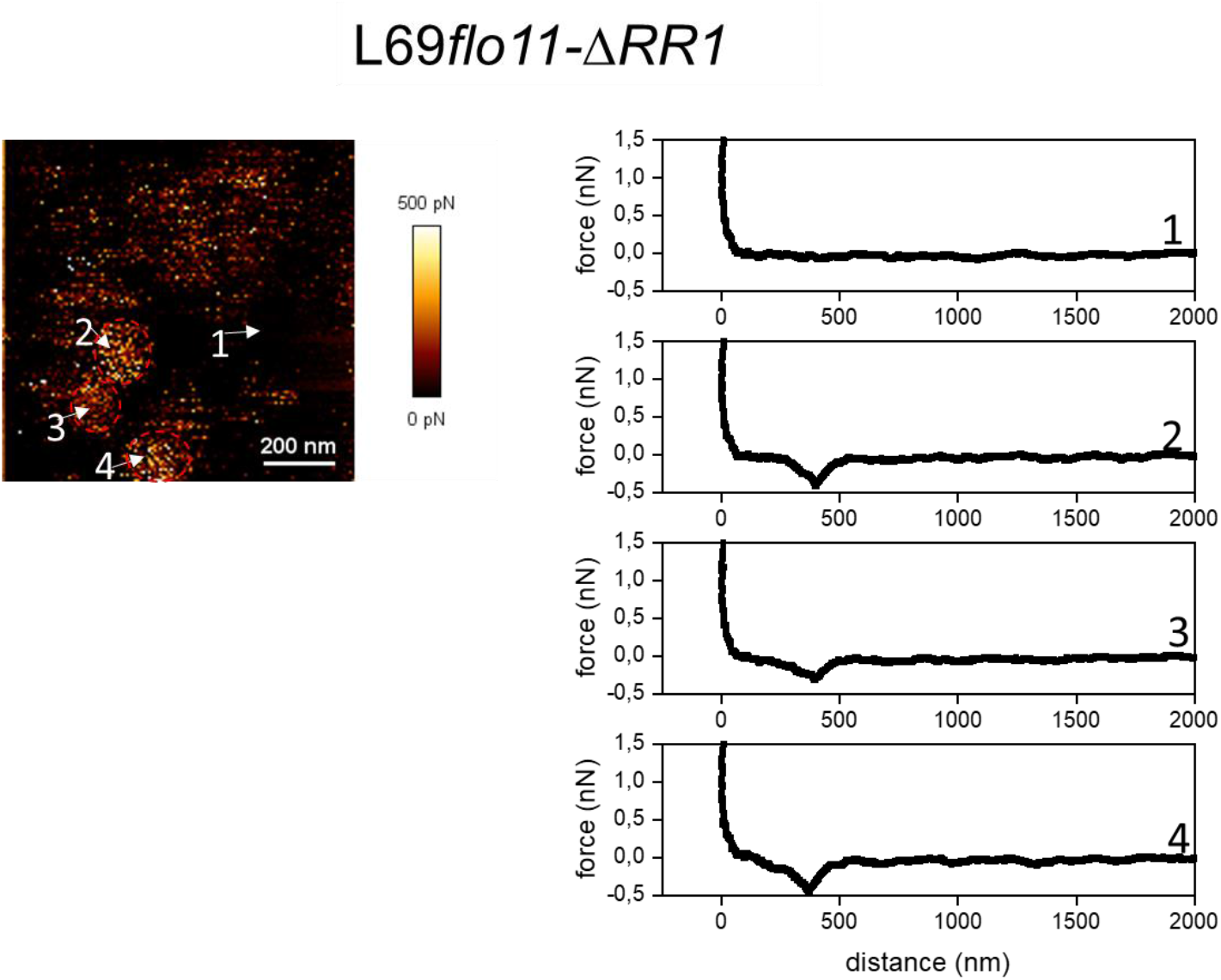
Cell surface analysis of L69*flo11-ΔRR1* using silicon-nitride (Si3N4) AFM cantilevers. Representative force-distance curves (right) at different location on the AFM adhesion image (left) indicated by a number obtained from a single cells of L69*flo11-ΔRR1* embedded in PDMS chamber.

**Figure S11:**
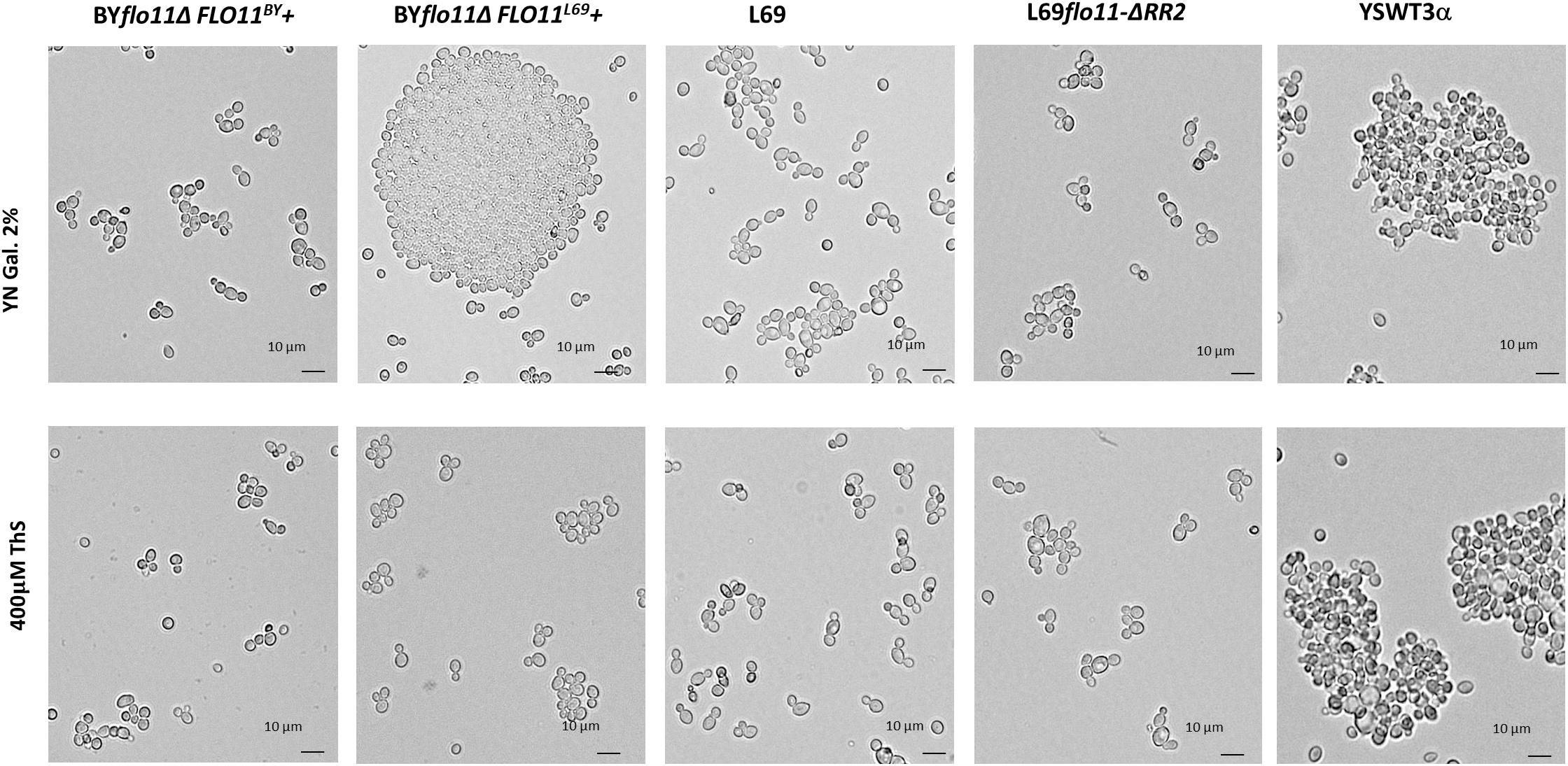
Effect of Thioflavine S on the Flollp-dependent cell-cell aggregation. BY4741 transformed with *FLO11*^*BY*^ or *FLO11*^*L69*^, L69, L69 expressing *FLO11* lacking RR2 *(L69flo11-Δ RR2)* and Σ1278b were cultivated in YN Gal. Cultures of these strains were harvested in exponential phase (O.D. around 1.0), and treated or not with 0.4 mM thioflavine S for 1h before microscopic observation.

**Figure S12:**
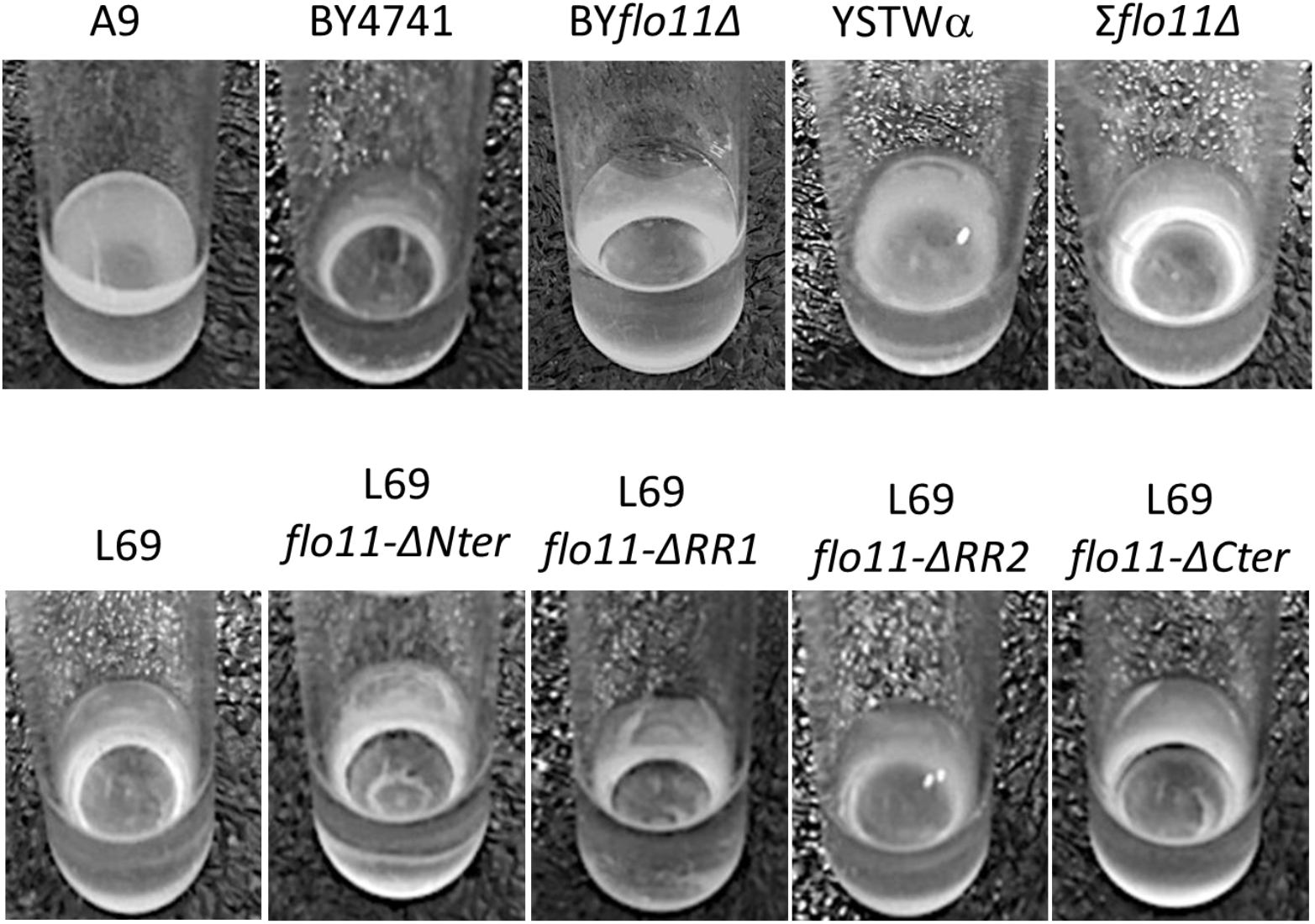
Assay of velum formation. The assay was carried out by static incubation for 7 days of yeast cells in flor medium at 23°C.

**Figure S13:**
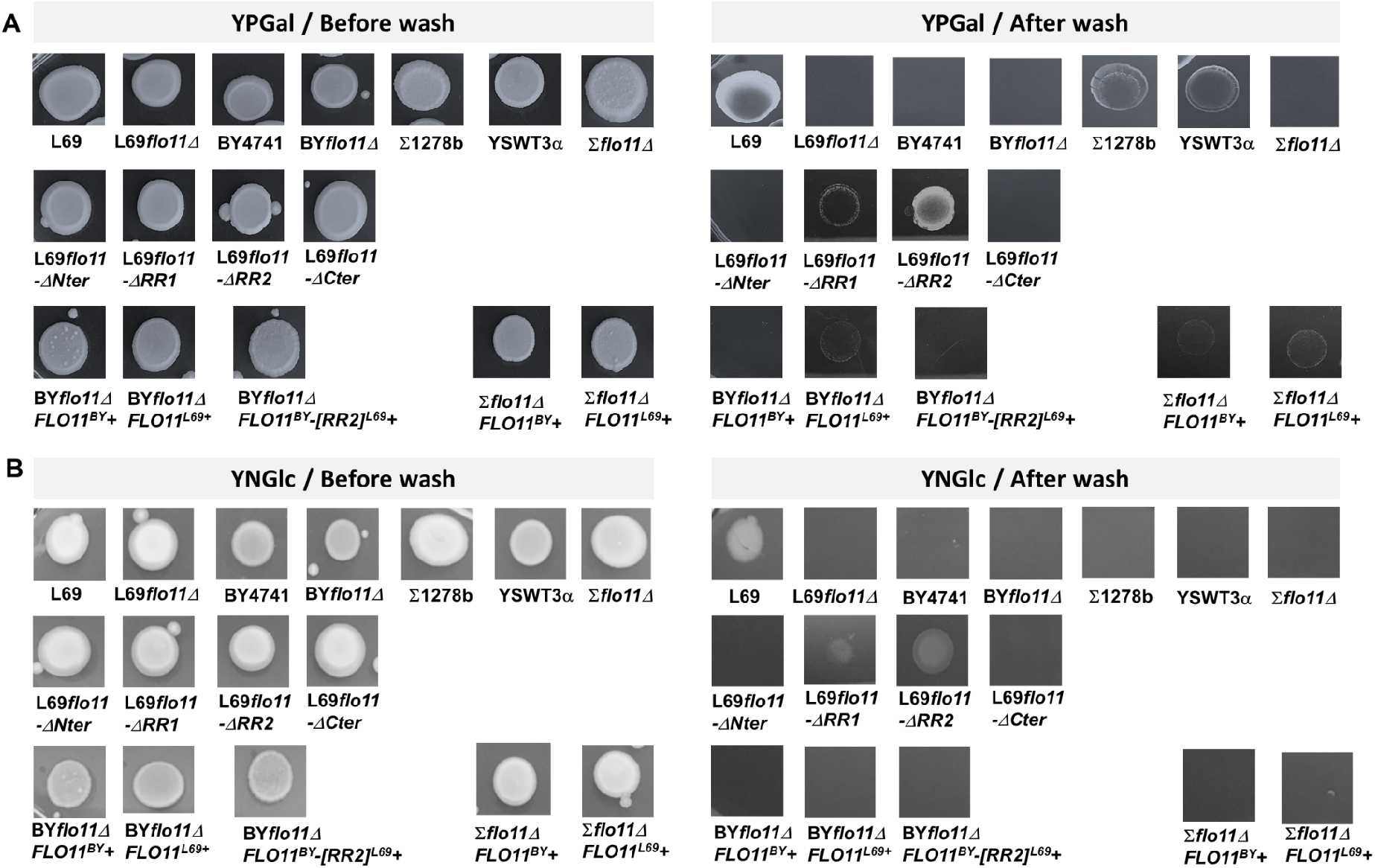
Invasive growth in agar by different yeast strains, and impact of the culture medium and of the domains of FLO11p on this phenotype. Yeast cells were initially cultivated in YNGal complemented with amino acids and uracil, except for BY*flo11Δ* and Σ*flo11Δ* expressing *FLO11*^*BY*^ or *FLO11*^*L69*^ on a pYES2.1 plasmid for which uracil was omitted. Cells were harvested at the entrance in stationary phase and deposited as patches on agar plates which were made with (A) rich galactose medium (YP Gal) or (B) with synthetic glucose medium complemented with amino acids and uracil, except for BY*flo11Δ* and Σ*flo11Δ* expressing *FLO11*^*BY*^ on *FLO11*^*L69*^ on a pYES2.1 plasmid for which uracil was omitted. Plates were incubated at 30°C for 8 days. They were then photographed before and after washing under a stream of water.

**Figure S14:**
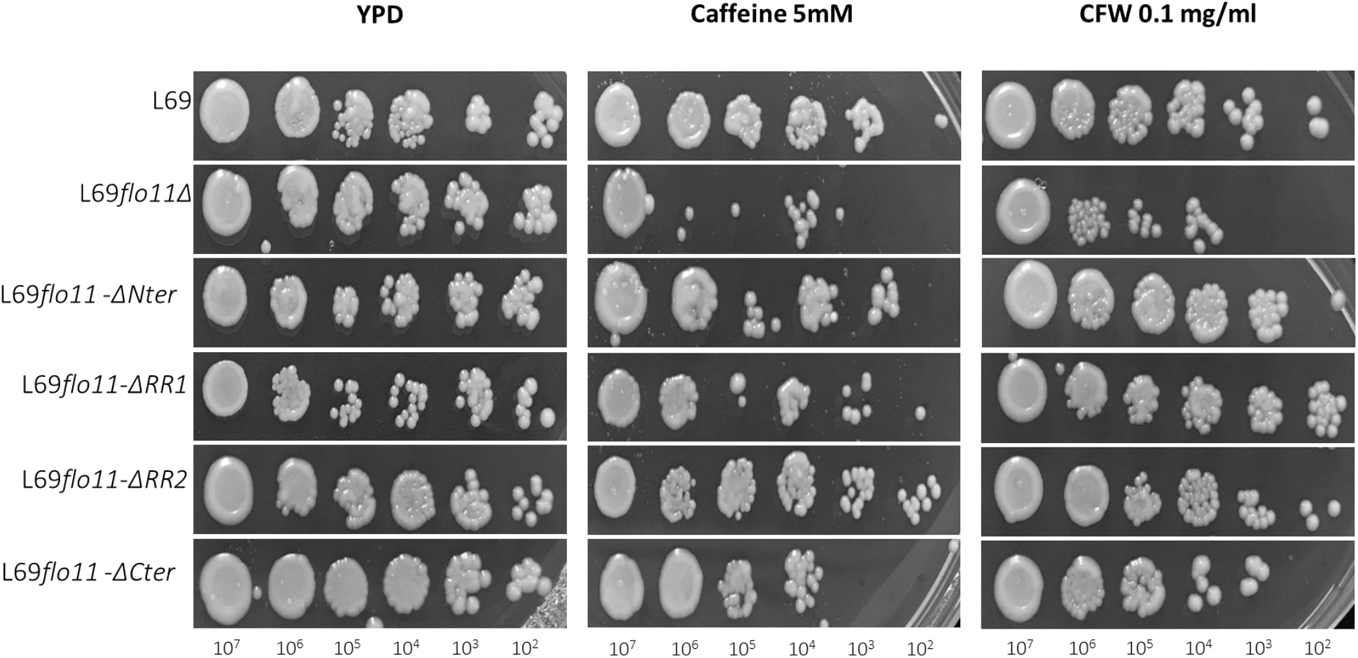
Sensitivity of L69 and L69 mutant strains to Caffeine and Calcofluor White (CFW) drugs. Cells were exponentially grown in YPD and were collected by centrifugation then resuspended in sterilize water at 10^8^ cells mL^-1^. Series of 10-fold dilutions were spotted on YPD agar plates in the absence or presence of CFW or Caffeine. Picture was taken after 2 days of growth at 30 °C.

**Table S1.** TANGO software analysis of β-aggregation motifs in Flo1, Flo5, Flo9, Flo10 and Flo11 protein from *Saccharomyces cerevisiae* S288c strain

| Flo1 |  |  | Flo5 |  |  | Flo9 |  |  | Flo10 |  |  | Flo11 |  |  |
| --- | --- | --- | --- | --- | --- | --- | --- | --- | --- | --- | --- | --- | --- | --- |
| AA position | motif | Mean $\beta$ aggregate (%) | AA position | motif | Mean $\beta$ aggregate (%) | AA position | motif | Mean $\beta$ aggregate (%) | AA position | motif | Mean $\beta$ aggregate (%) | AA position | motif | Mean $\beta$ aggregate (%) |
| 7-21 | YMFLAVFTLLAL<br>TSV | 82.2 | 9-22 | IFLVILAFLALINVA | 82 | 6-22 | YCLLLAIVTLLG<br>LTNVV | 77.55 | 7-23 | YIFLTGLFLLSVANV<br>AL | 81.7 | 5-16 | FLLAYLVLSLLF | 96 |
| 308-312 | TVIVI | 87.9 | 207-215 | TVYMYAGYY | 30.5 | 118-122 | IIAYW | 72 | 1028-1035 | VVTVYSTW | 48.5 | 1033-1042 | VTTVVSTTVV | 75.8 |
| 353-357 | TIIVI | 87.3 | 308-312 | TVIVI | 73.2 | 207-215 | TVYMYAGFY | 50.9 | 1115-1119 | VLISV | 47 | 1056-1061 | ITTTFV | 56 |
| 398-402 | TIIVI | 87.3 | 353-357 | TVIVI | 87.8 | 308-312 | TVIVI | 87.9 | 1157-1168 | ISIFIASLLAI | 89.8 | 1133-1144 | TLVTTAVTTTVV | 84.8 |
| 443-447 | TIIVI | 87.3 | 398-402 | TVIVI | 87.8 | 353-357 | TIIVI | 87.3 |  |  |  | 1356-1362 | FMWLLLA | 85.3 |
| 488-492 | TIIVI | 87.3 | 443-447 | TVIVI | 87.8 | 398-402 | TIIVI | 87.3 |  |  |  |  |  |  |
| 533-537 | TIIVI | 87.2 | 488-492 | TVIVI | 87.8 | 443-447 | TIIVI | 87.3 |  |  |  |  |  |  |
| 578-562 | TIIVI | 87.2 | 533-537 | TVIVI | 87.9 | 498-492 | TIIVI | 87.3 |  |  |  |  |  |  |
| 623-627 | TIIVI | 87.2 | 578-582 | TVIVI | 87.9 | 533-537 | TIIVI | 87.2 |  |  |  |  |  |  |
| 667-672 | TIIVI | 87.2 | 623-627 | TVIVI | 87.9 | 578-582 | TIIVI | 87.2 |  |  |  |  |  |  |
| 713-717 | TIIVI | 87.2 | 783-788 | TLVTVT | 31.7 | 623-627 | TIIVI | 87.2 |  |  |  |  |  |  |
| 758-762 | TVIVI | 87.8 | 802-811 | AIVSTATVTV | 45.2 | 668-672 | TIIVI | 87.2 |  |  |  |  |  |  |
| 803-807 | TVIVI | 87.8 | 855-859 | TVVTI | 37.2 | 713-717 | TIIVI | 87.2 |  |  |  |  |  |  |
| 848-852 | TVIVI | 87.8 | 906-911 | TLVTVT | 36.1 | 758-762 | TVIVI | 87.8 |  |  |  |  |  |  |
| 893-897 | TVIVI | 87.8 | 1063-1074 | LSVFIASLLAI | 87 | 803-807 | TVIVI | 87.8 |  |  |  |  |  |  |
| 938-942 | TVIVI | 87.8 |  |  |  | 848-852 | TVIIV | 87.7 |  |  |  |  |  |  |
| 983-987 | TVIVI | 87.8 |  |  |  | 1021-1026 | TLVTVT | 31.9 |  |  |  |  |  |  |
| 1028-1032 | TVIVI | 87.9 |  |  |  | 1040-1049 | AIVSTATVTV | 42.6 |  |  |  |  |  |  |
| 1073-1077 | TVIVV | 88.5 |  |  |  | 1093-1097 | TVVTI | 37.4 |  |  |  |  |  |  |
| 1234-1239 | TLVTVT | 31.7 |  |  |  | 1144-1149 | TLVTVT | 37.7 |  |  |  |  |  |  |
| 1254-1262 | IVSTATVTV | 45.9 |  |  |  | 1175-1182 | VVTVYSTW | 82.6 |  |  |  |  |  |  |
| 1299-1303 | TVVTI | 37.2 |  |  |  | 1310-1321 | LSVFIASLLAI | 87 |  |  |  |  |  |  |
| 1350-1355 | TLVTVT | 36 |  |  |  |  |  |  |  |  |  |  |  |  |
| 1525-1536 | LSVFIASLLAI | 87 |  |  |  |  |  |  |  |  |  |  |  |  |

**Table S2.**
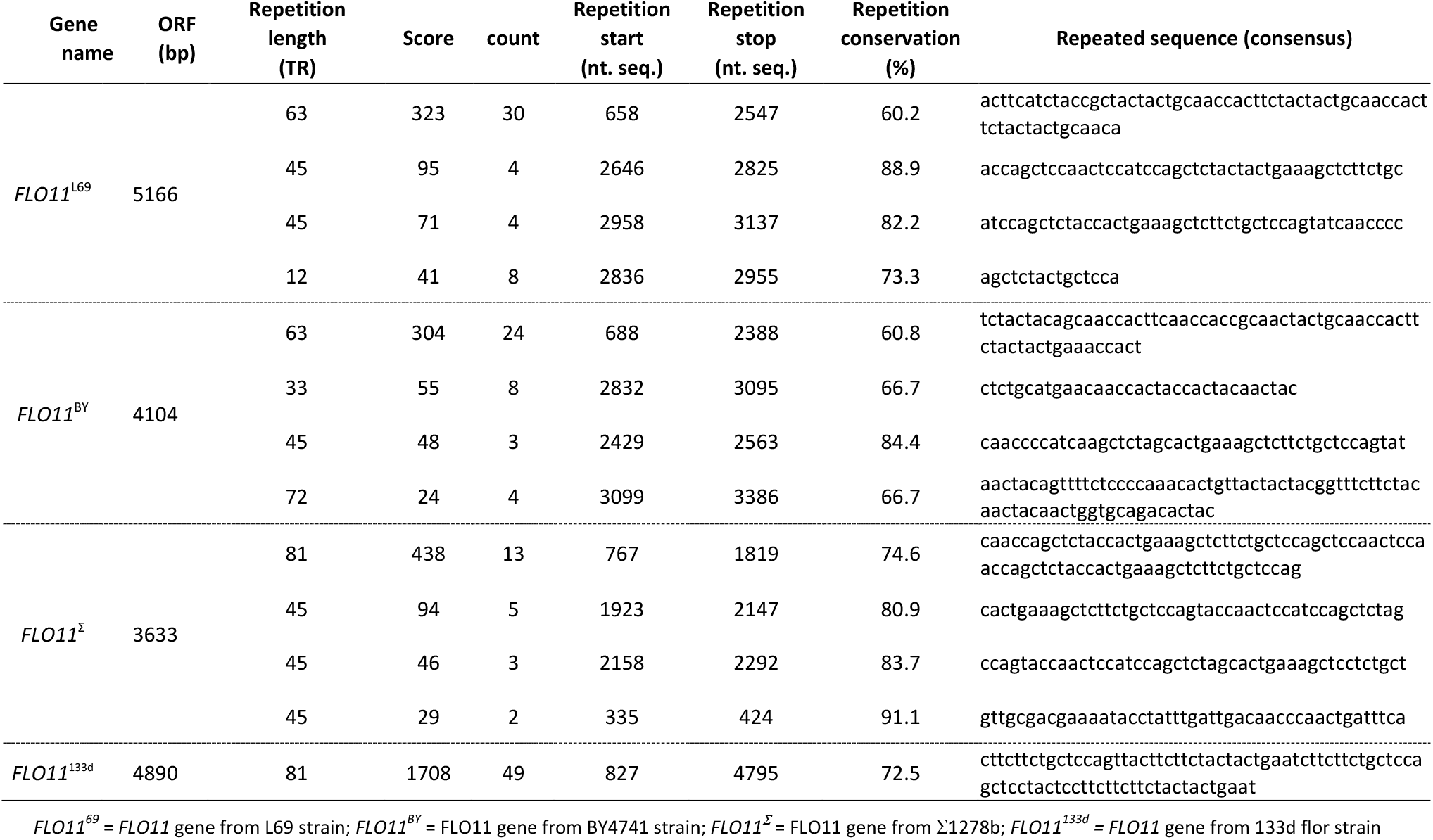
Search for intragenic repeats using EMBOSS ETANDEM software

**Table S3.**
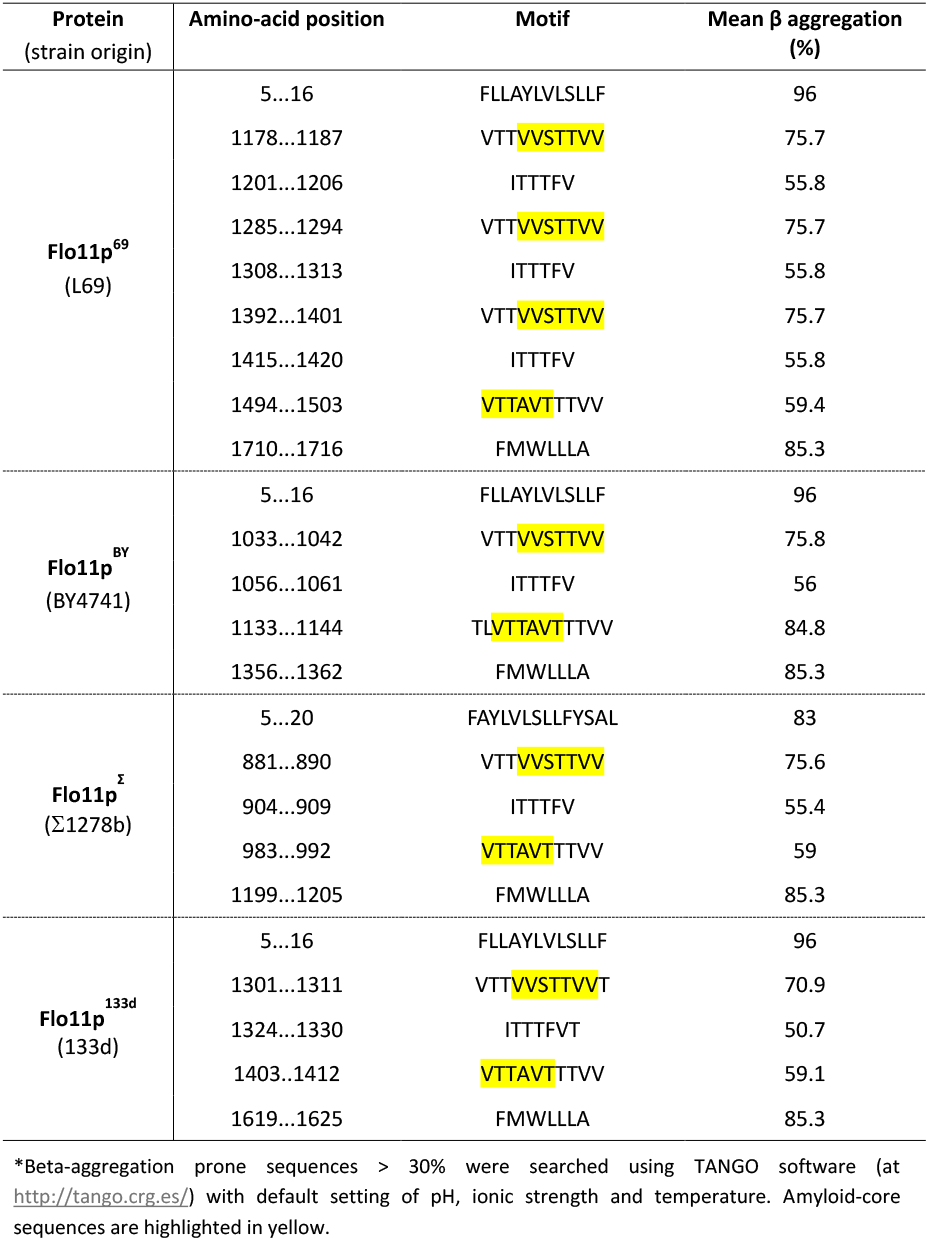
Search for β-aggregation prone sequence in the different Flo11 proteins using TANGO software (http://tango.crg.es/)

**Table S4.** Yeast strains used or constructed in this study.

| Strain | Genotype/Remarks | Source |
| --- | --- | --- |
| L69 | *Saccharomyces cerevisiae* prototrophic diploid. | Lallemand Inc. |
| L69*flo11Δ* | L69 *flo11::KanMX4*. | This study |
| L69*flo11Δ-Nter* | L69 diploid strain expressing *FLO11* deleted of its N-terminus domain. | This study |
| L69*flo11Δ-RR1* | L69 diploid strain expressing *FLO11* deleted of the intragenic RR1. | This study |
| L69*flo11Δ-RR2* | L69 diploid strain expressing *FLO11* deleted of the RR2 intragenic domain | This study |
| L69*flo11Δ-Cter* | L69 diploid strain expressing *FLO11* deleted of its C-terminal domain | This study |
| BY4741 | MATa *his3Δ1 leu2Δ0 met15Δ0 ura3Δ0*. | Euroscarf |
| BY*flo11Δ* | BY4741 *flo11::KanMX4*. | YKO Open biosystem |
| Σ1278b | Prototrophic diploid strain | Lab source |
| YSWT3α | MATα *can1Δ::Ste2pr-spHIS5 lyp1Δ::Ste3pr-LEU2 his3::hisG leu2Δ0 ura3Δ0*. Derived from Σ1278b | C.Boone (Univ. Toronto) |
| Σ*flo11Δ* | YSWT3α *flo11 ::kanMX4* | C.Boone (Univ. Toronto) |
| BY*flo11Δ FLO11*<sup>BY+</sup> | BY4741 *flo11::KanMX4* transformed with pYES *FLO11*<sup>BY</sup>. | This study |
| BY*flo11Δ FLO11*<sup>L69+</sup> | BY4741 *flo11::KanMX4* transformed with pYES-*FLO11*<sup>L69</sup> | This study |
| BY*flo11ΔFLO11*<sup>BY</sup>[RR2]<sup>L69+</sup> | BY4741 *flo11::KanMX4* haploid strain transformed with pYES-*FLO11*[RR2]<sup>L69</sup> | This study |
| Σ*flo11Δ FLO11*<sup>BY+</sup> | YSWT3α *flo11 ::kanMX4* strain transformed with pYES-*FLO11*<sup>BY</sup> | This study |
| Σ*flo11Δ FLO11*<sup>L69+</sup> | YSWT3α *flo11 ::kanMX4* strain transformed with pYES-*FLO11*<sup>L69</sup> p | This study |
| BY4741+pFLO8 | BY4741 transformed with pGP564_FLO8 vector. | This study |
| A9 | MATa/MATHO/HO | M. Budroni (Univ di Sassari, Italy) |

**Table S5:** Plasmids constructed in this work

| Plasmids | description | Source |
| --- | --- | --- |
| pYES2.1 TOPO TA | E.coli /yeast shuttle vector with His Tag (6x), V5 Epitope Tag, Amp<sup>R</sup>, *GAL1* promotor, *URA3* marker. | Life Technologies |
| pYES-*FLO11*<sup>L69</sup> | Cloning of *FLO11*<sup>L69</sup> into pYES2.1 TOPO vector. | This study |
| pYES-*FLO11*<sup>BY</sup> | Cloning of *FLO11*<sup>BY</sup> into pYES2.1 TOPO vector. | This study |
| pYES-*FLO11*<sup>BY</sup>-[*RR2*]<sup>L69</sup> | Cloning of *FLO11*<sup>BY</sup>[*RR2*]<sup>L69</sup> chimeric gene in pYES2.1 | This study |
| pGP564_*FLO8* | pGP564 vector carrying GLE2, YER107W-A, *FLO8*, KAP123 and SWI4 | Open Biosystem |

**Table S6.**
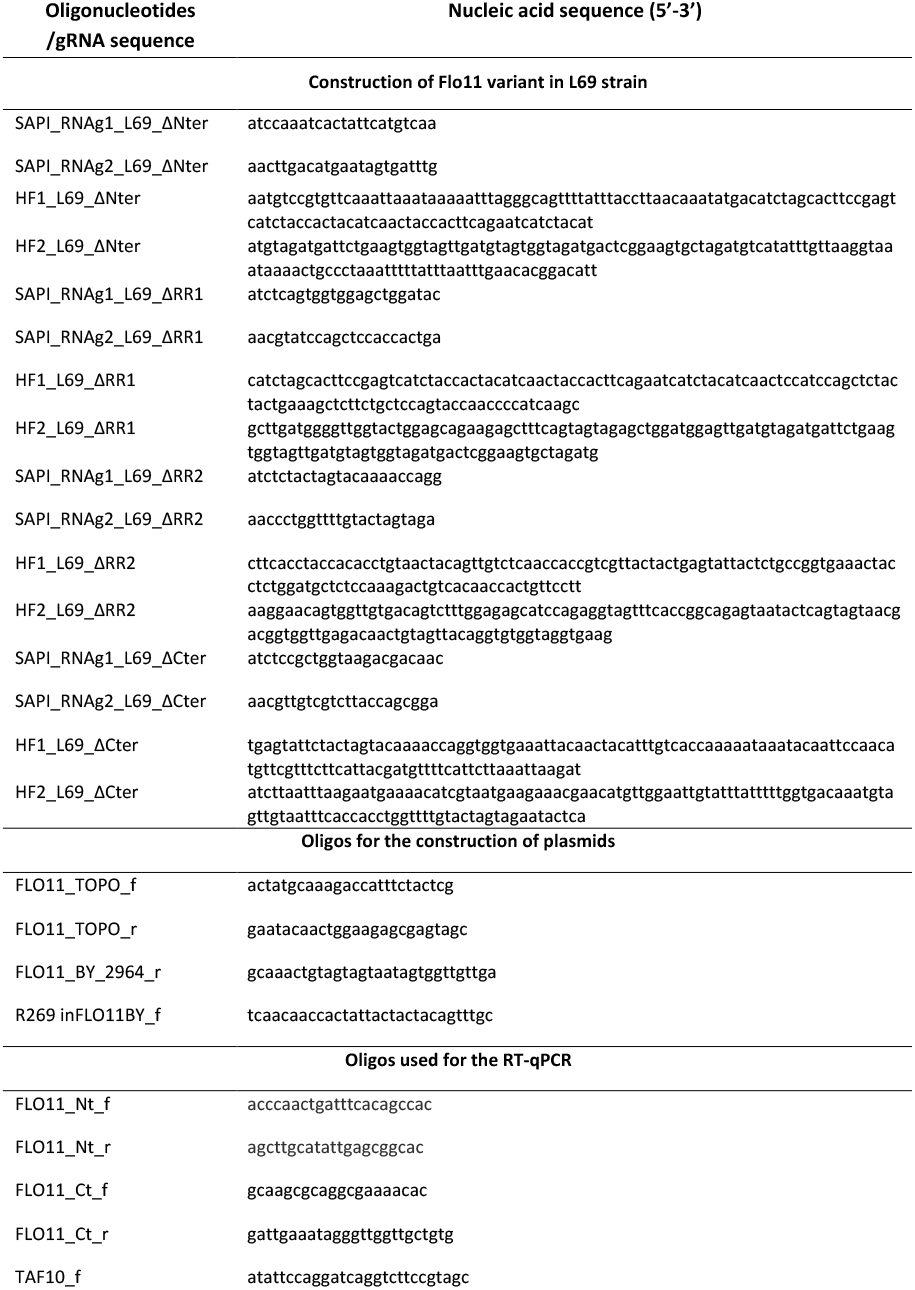
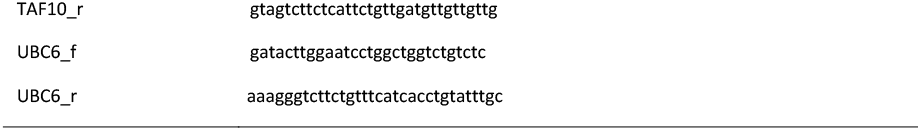
Oligonucleotides used in this study.

